# Silversol^®^ (a colloidal nanosilver formulation) inhibits growth of antibiotic-resistant *Staphylococcus aureus* by disrupting its physiology in multiple ways

**DOI:** 10.1101/2024.04.23.590707

**Authors:** Nidhi Thakkar, Gemini Gajera, Chhaya Godse, Anselm DeSouza, Dilip Mehta, Vijay Kothari

## Abstract

Antibacterial effect of a colloidal nanosilver formulation Silversol^®^ was investigated against an antibiotic-resistant strain of *Staphylococcus aureus*. Lower concentrations of the test formulation exerted bacteriostatic and, its higher concentrations exerted bactericidal effect against this pathogen. Silversol^®^ at sub-lethal concentration was found to disturb multiple physiological traits of *S. aureus* such as growth, antibiotic susceptibility, membrane permeability, efflux, protein synthesis and export, biofilm and exopolysaccharide production, etc. Transcriptome data revealed the genes coding for transcriptional regulators, efflux machinery, transferases, β-lactam resistance, oxidoreductases, metal homeostasis, virulence factors, and arginine biosynthesis to get expressed differently under influence of the test formulation. Genes (*argG* and *argH*) involved in arginine biosynthesis emerged among the major targets of Silversol^®^ in *S. aureus*.

## 1. Introduction

Historically, silver has long been in use for various therapeutic purposes. It is an important ingredient of multiple traditional medicine formulations (Saper et al., 2004; Inder and Kumar, 2014; Parimalam et al., 2020). Modern times has also witnessed numerous reports (Burange et al., 2021; Korkmaz, 2020; Khan et al., 2023) describing different types of biological activities of silver. Antimicrobial activity of different forms of silver has also been documented e.g., antiviral (Balasubramaniam et al., 2020; Jain et al., 2021; Ratan et al., 2021), antibacterial (Balasubramaniam et al., 2020; Ibrahim et al., 2020; Bruna et al., 2021), antifungal (Mussin et al., 2022; Matras et al., 2022), antiprotozoal (Abou Elez et al., 2023), and anthelmintic (Majumdar and Kar, 2023; Gajera et al., 2023a). Different forms of silver (e.g., metallic, colloidal, silver salts) may have different mechanisms of action, and varying magnitude of biological activity. Antibacterial activity of silver assumes importance in face of the AMR (antimicrobial resistance) pandemic. Literature review does not provide strong indications on bacteria developing resistance against silver (Wu et al., 2022). Silver is reported to exert its bactericidal activity by modulating cell membrane permeability, triggering generation of reactive oxygen species (ROS), and interrupting DNA replication (Ahmad et al., 2017; Mazur et al., 2020; Reeves et al., 2020; Yin et al., 2020; Jiang et al., 2022). However, investigation is still warranted for elucidating additional details on molecular mechanisms associated with antibacterial activity of silver.

Since metallic form of silver is believed to be more potent than its ionic form (de Souza et al., 2021), we chose a solution of colloidal silver for the purpose of this study. Target pathogen for this study was an antibiotic-resistant strain of *Staphylococcus aureus*. This bacterium is one of the most notorious and versatile bacterial pathogens, whose antibiotic-resistant strains have been listed by Centers for Disease Control and Prevention (CDC; https://www.cdc.gov/drugresistance/biggest-threats.html), World Health Organization (WHO; https://www.who.int/publications/i/item/WHO-EMP-IAU-2017.12), and Department of Biotechnology of the Indian government (DBT; https://dbtindia.gov.in/sites/default/files/IPPL_final.pdf) among priority pathogens, against whom there is a pressing need to discover new antibacterial agents (Weiner et al., 2016). This study investigated effect of Silversol^®^ (a colloidal nano-silver formulation) on *S. aureus*’s growth, various important phenotypic traits, and gene expression profile at the whole transcriptome level. Silversol is already in clinical use for multiple therapeutic purposes e.g., management of oral health (Tran et al., 2019), skin diseases, burns, diabetic ulcer, and wound-care/ wound-disinfection (https://www.rxsilver.com/index_htm_files/ABLSilversafety.pdf). *S. aureus* being one of the common clinical isolates (Reyes et al., 2023; Phan et al., 2023), we attempted to gain insight into Silversol’s antibacterial activity against this pathogen.

## 2. Methods

### 2.1. Colloidal silver formulation

The Silversol^®^ test formulation (100 ppm; batch number: V-Silwater22-47), originally formulated by American Biotech Labs (USA), was sourced from Viridis BioPharma Pvt. Ltd., Mumbai, India. This colloidal silver mixture is documented to possess multiple biological effects (de Souza et al., 2021). It comprises of zero-valent metallic silver particles in their elemental form that are covered in silver oxide. The manufacturer claims its particle size range to lie between 5 and 50 nm.

### 2.2. Bacterial culture

The strain of *S. aureus* utilized in this study (ATCC 43300) was procured from HiMedia, Mumbai. Though this strain is claimed to be resistant to methicillin and oxacillin [https://www.atcc.org/products/43300], disc diffusion assay in our lab did yield a zone of inhibition of 26.5±9.19 and 23±7.07 mm surrounding methicillin and oxacillin discs (Table S1 and Figure S1). However, our transcriptome of this strain, conducted as a part of this study, showed this strain to be positive for expression of *mecA* gene, whose presence is considered a defining character for labelling *S. aureus* as methicillin-resistant (EUCAST guidelines, 2017). Antibiogram of this strain generated in our lab showed it to be resistant to four macrolide antibiotics (erythromycin, clindamycin, azithromycin, and clarithromycin), and intermediate to amoxycillin/clavulanic acid. For characterizing the test strain in terms of virulence, we challenged the nematode worm *Caenorhabditis elegans* with it (OD_764_=1.50), and it was able to kill 100% worms within 24 h (Figure S2). In summary, the *S. aureus* strain used in this study can be said to possess the traits of antibiotic-resistance and virulence.

This bacterium was maintained on Tryptone soy agar (HiMedia) slants. For the purpose of different assays described in this report, (unless described otherwise) it was grown in tryptone soy broth (pH 7.0±0.2; HiMedia) for 24±1 h at 35 °C. Inoculum was standardized as OD_625_ between 0.08-0.10 (at par to McFarland turbidity standard 0.5).

### 2.3. Growth inhibition assay

Effect of Silversol on bacterial growth was quantified through broth dilution assay. *S. aureus* was inoculated (at 10%v/v) in tryptone soy broth with or without Silversol (0.1-60 ppm), followed by incubation at 35 °C for 24±1 h. Total volume of the system was 2 mL in 15 mL test tubes. Random intermittent shaking was provided during incubation. At the end of incubation, cell density was measured at 764 nm (Agilent Cary 60 UV-Vis).

The lowest concentration of Silversol capable of inhibiting ≥80% growth was taken as the minimum inhibitory concentration (MIC) (Turnidge, 2015). From each tube showing visible absence of growth, 0.1 mL of culture broth was spread on tryptone soy agar plates. These agar plates were observed for the appearance of bacterial growth over an incubation (at 35°C) period of 72 h for the determination of the minimum bactericidal concentration (MBC). Extended incubation till 72 h was made to differentiate true bactericidal effect from any possible post-antibiotic effect (Ramadan et al., 1995; Pfaller et al., 2004). The lowest concentration of Silversol, which could inhibit the appearance of growth on agar plates almost completely, was taken as MBC. Vancomycin was used as a positive control. Growth curve of the test bacterium in absence and presence of Silversol (at sub-MIC level) was also generated, wherein incubation was carried out under continuous shaking (120 rpm).

### 2.4. Antibiotic susceptibility assay

Antibiogram of Silversol pre-exposed-*S. aureus* was generated through disc diffusion assay performed in accordance to the Clinical and Laboratory Standards Institute (CLSI) guidelines (Simner et al., 2022), and was compared with that of control culture (which received no Silversol pre-exposure). *S. aureus* was grown in tryptone soy broth, with or without Silversol. Following overnight incubation (24±1 h), 1 mL culture broth was used for cell density quantification at 764 nm, and cells from the remaining 1 mL culture were pelleted down through centrifugation (10 min at 13,600 g). The resulting pellet was then resuspended in 1 mL phosphate buffer (HiMedia; pH 7.0 ± 0.2), followed by centrifugation. The resulting cell pellet was utilized to prepare the inoculum for the subsequent disc diffusion assay by resuspending it in normal saline. OD_625_ of the inoculum was adjusted to range from 0.08-0.10, to achieve equivalence to McFarland turbidity standard 0.5. One hundred µL of this inoculum was then spread onto cation-adjusted Muller-Hinton agar (HiMedia) in 150 mm plates (Borosil), followed by placing antibiotic discs (Icosa G-I PLUS; HiMedia, Mumbai) onto the agar surface. The plates were then incubated at 35°C for 18±1 h, followed by the observation and measurement of the diameter of zone of inhibition.

### 2.5. Biofilm assay

Biofilm formation is an important virulence trait, and hence effect of Silversol on biofilm forming ability of *S. aureus*, as well as on pre-formed biofilm was investigated through four different biofilm assays using methodology described in our previous publication (Gajera et al., 2023b). A flow diagram depicting all the biofilm assays is provided in Figure 1, wherein biofilm quantification was achieved through crystal violet assay (Hirshfield et al., 2009), and biofilm viability was assessed through MTT assay (Trafny et al.,2013). Vancomycin at sub-MIC level (3 ppm) was used as a positive control.

**Figure 1.**
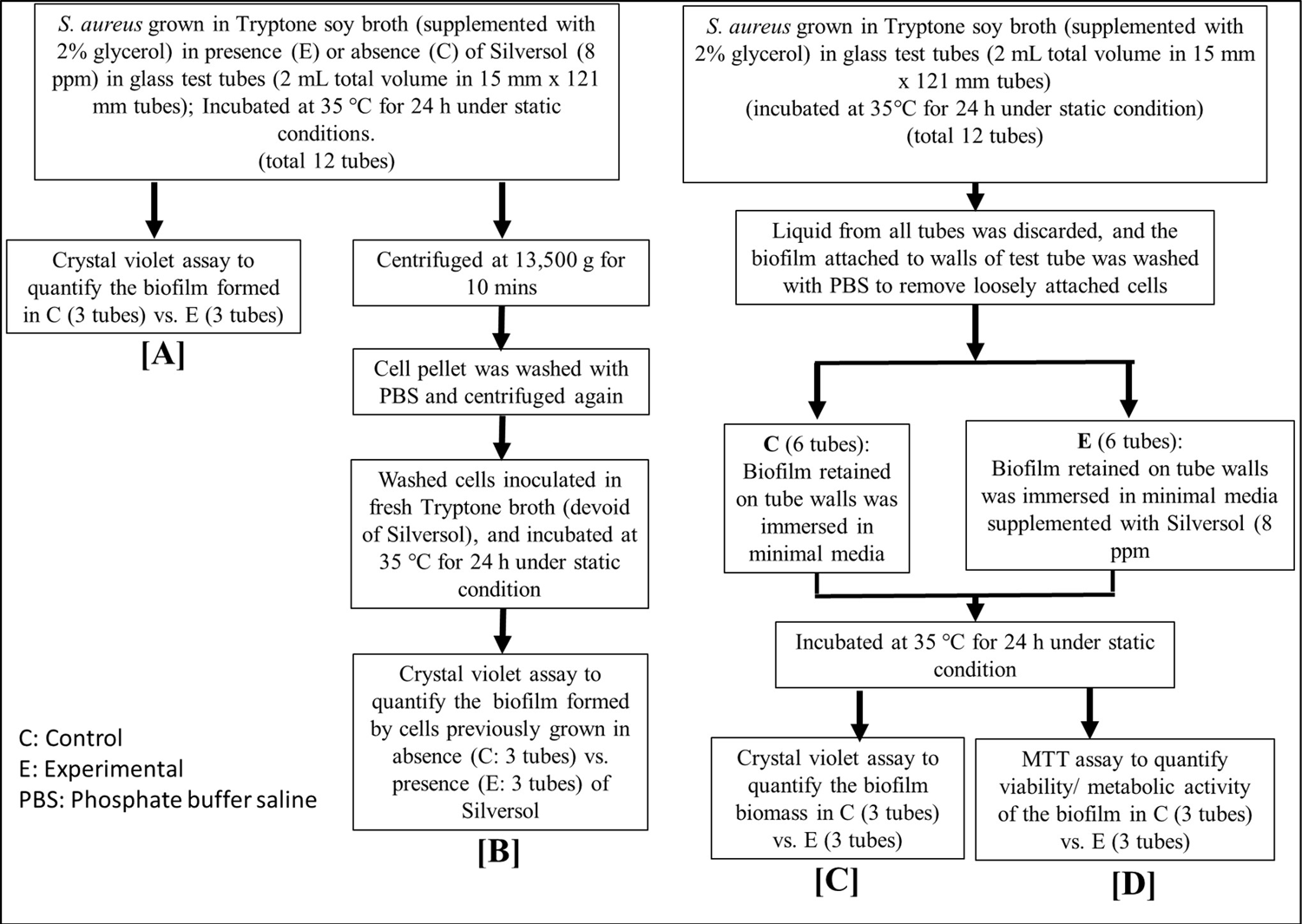
Flowchart depicting schematic of all biofilm assays **(A)** Quantification of biofilm formation in presence or absence of Silversol; **(B)** Quantification of biofilm formation by silver-pre-treated vs. non-pre-treated *S. aureus* cells; **(C)** Quantification of biofilm eradication after adding Silversol onto pre-formed biofilm; **(D)** Quantification of Silversol’s effect on metabolic activity of pre-formed biofilm

For the crystal violet assay, the biofilm containing tubes (after discarding the inside liquid) were washed with phosphate buffer saline (PBS) so as to remove all non-adherent (planktonic) bacteria, and air-dried for 15 min. Then, each of the washed tubes was subjected to staining with 2 mL of 0.4% aqueous crystal violet (Central Drug House, Delhi) solution for 30 min. Each tube was then washed twice with two mL of sterile distilled water and immediately de-stained with 2 mL of ethanol (95%). After 45 min of de-staining, one mL of destaining solution was transferred into separate tubes, and absorbance was quantified at 580 nm (Agilent Cary 60 UV-Vis).

For the MTT assay, the biofilm-containing tubes (after discarding the inside liquid) were washed with PBS so as to remove all non-adherent (planktonic) bacteria, and air-dried for 15 min. Then 1.8 mL of minimal media (MgSO_4_ 0.1 g/L, K_2_HPO_4_ 5 g/L, NH_4_Cl 2 g/L, sucrose 15 g/L, NaCl 1 g/L, yeast extract 0.1 g/L, pH 7.4 ± 0.2) was added into each tube, followed by addition of 200 µL of 0.3% MTT [3-(4,5-Dimethylthiazol-2-yl)-2,5-iphenyltetrazolium Bromide; HiMedia]. Then after incubation at 35°C for 24 h, all liquid content was discarded, and the remaining purple formazan derivatives were dissolved in two mL of DMSO and absorbance was measured at 540 nm.

### 2.6. Exopolysaccharide (EPS) quantification

*S. aureus* was grown in 100 mL flasks containing 20 mL of tryptone soy broth (containing or not containing Silversol). Incubation was provided at 35°C for 24±1 h under static condition with intermittent shaking. At the end of incubation, cell density was quantified as OD_764_. For EPS quantification (Andhare et al., 2017), culture broth was subjected to centrifugation (13,600 g for 10 min). To the resulting supernatant, chilled acetone (Merck) was added in 1:2 ratio, and allowed to stand for 30 min for the EPS to get precipitated. Thus, precipitated EPS was then separated by filtration through pre-weighed Whatman # 1 filter paper (Whatman International Ltd., England). Filter paper containing EPS on it was then dried for 24 h, and weight of EPS on the paper was calculated.

### 2.7. Efflux assay

Ethidium bromide (EtBr) efflux assay with *S. aureus* was performed using the method described in Mullin et al. (2004). *S. aureus* cells, grown overnight in tryptone soy broth medium, were loaded with 8 ppm of EtBr (HiMedia) in the presence of 75 ppm reserpine (HiMedia; positive control) or Silversol (8 ppm). Any effective efflux inhibitor like reserpine will inhibit efflux during this loading step. Cells were incubated at 35°C for 20 min and then pelleted by centrifugation (13,600 g; 10 min). The medium was decanted, and the cell pellet was resuspended in fresh tryptone soy broth medium, either with or without Silversol (8 ppm), to an optical density OD_764_ = 0.20. EtBr efflux was then determined by quantification of fluorescence at excitation and emission wavelengths of 530 and 600 nm in spectrofluorometer (JASCO FP-6500).

### 2.8. Protein estimation

Extracellular protein present in bacterial culture (grown in presence or absence of Silversol) supernatant, and intracellular protein in the cell lysate was quantified through Folin-Lowry method (Lowry et al., 1951; Dulley and Grieve, 1975). After measuring cell density, one mL of *S. aureus* culture was centrifuged (13,600 g), and the resulting supernatant was used for extracellular protein estimation. The remaining cell pellet was subjected to lysis (Mishra et al., 2019) for release of intracellular proteins. Briefly, the cell pellet was washed with phosphate buffer (pH 7.4), and centrifuged (13,600 g). Resulting pellet was resuspended in 1 mL of chilled lysis buffer (0.876 g NaCl, 1 mL of Triton X 100, 0.5 g sodium deoxycholate, 0.1 g sodium dodecyl sulphate, and 0.60 g Tris HCl, in 99 mL of distilled water), and centrifuged (500 rpm) for 30 min at 4°C for agitation purpose. This was followed by further centrifugation (13,600 g at 4°C) for 20 min. Resulting cell lysate (supernatant) was used for protein estimation.

### 2.9. Membrane permeability assay

Propidium Iodide (PI) uptake assay (Crowley et al., 2016; Mombeshora and Mukanganyama, 2019) was performed to quantify membrane permeability. *S. aureus* cells grown overnight in tryptone soy broth were centrifuged, and cell pellet was dissolved in PBS to attain OD_764_ = 1.50±0.05. This cell suspension was exposed to Silversol (8 ppm) for 30 min at 35 °C in an incubator. The negative control contained cells with no Silversol-exposure. After incubation, two ml of each test sample was centrifuged at 13,600 g. The resulting pellet was washed with PBS, resuspended in PBS, and then PI (Invitrogen) at a final concentration of 10 ppm was added into the suspension. This mixture was kept in the dark for 10 min, followed by quantification of fluorescence, employing 544 nm as excitation and 612 as emission wavelength, using spectrofluorometer (JASCO FP-6500). Triton-X (final concentration 0.5%v/v) was kept as positive control.

### 2.10. Transcriptome analysis

To gain insights into the molecular mechanisms through which silver could inhibit bacterial growth and modulate various physiological traits, gene expression profile of *S. aureus* challenged with sub-MIC of Silversol (8 ppm) was compared with that of control culture at the whole transcriptome level. Overall workflow of this whole transcriptome analysis (WTA) aimed at obtaining a holistic picture regarding mode of action of this formulation is presented in Figure S3.

#### 2.10.1. RNA extraction

RNA from bacterial cells was extracted by Trizol (Invitrogen Bioservices; 343909) method. Precipitation was done by isopropanol followed by washing with 75% ethanol, and RNA was dissolved in nuclease free water. Extracted RNA was quantified on Qubit 4.0 fluorometer (Thermofisher; Q33238) using RNA HS assay kit (Thermofisher; Q32851) following manufacturer’s protocol. Purity and concentration of RNA was measured on Nanodrop 1000. Finally, to obtain RIN (RNA Integrity Number) values, RNA was checked on the TapeStation using HS RNA ScreenTape (Agilent) (Table S2).

#### 2.10.2. Library preparation

Final libraries were quantified through Qubit 4.0 fluorometer (Thermofisher; Q33238) using DNA HS assay kit (Thermofisher; Q32851) following manufacturer’s protocol. To identify the insert size of the library, it was queried on Tapestation 4150 (Agilent) employing high sensitive D1000 screentapes (Agilent; 5067-5582) as per manufacturers’ protocol. Acquired sizes of all libraries are reported in Table S2.

#### 2.10.3. Genome annotation and functional analysis

Quality assessment of the raw fastq reads of the sample was performed using FastQC v.0.11.9 (default parameters) (Andrews, 2010). The raw fastq reads were pre-processed using Fastp v.0.20.1 (Chen et al., 2018), followed by quality reassessment using FastQC.

The *S. aureus* genome (GCA_000006765.1_ASM676v1) was indexed using bowtie2-build (Langmead and Salzberg, 2012) v2.4.2 (default parameters). The processed reads were mapped to the indexed *S. aureus* genome using bowtie2 v2.4.2 parameters. The aligned reads from the individual samples were quantified using feature count v. 0.46. 1 (Liao et al., 2014) to obtain gene counts. These gene counts were used as inputs to edgeR (Robinson et al., 2010) using exact test (parameters: dispersion 0.1) for differential expression estimation. The up- and down-regulated sequences were extracted from the *S. aureus* coding file and subjected to blast2go (Conesa et al., 2008) for annotation to extract the Gene Ontology (GO) terms. These GO terms were subjected to the wego (Ye et al., 2018) tool to obtain gene ontology bar plots. All the raw sequence data has been submitted to the Sequence Read Archive. The relevant accession number is SRX15156375 (https://www.ncbi.nlm.nih.gov/sra/SRX15156375).

### 2.11. Network analysis

From the list of differentially expressed genes (DEG) in Silversol-exposed *S. aureus*, those satisfying dual filter criteria of log fold-change ≥2 and False Discovery Rate (FDR) ≤0.05 were chosen for further network analysis. The list of such DEG was fed into the STRING (v.11.5) database (Szklarczyk et al., 2019) to generate the PPI (Protein-Protein Interaction) network. Members of this PPI network were then arranged in decreasing order of ‘node degree’ (a measure of connectivity with other genes or proteins), and those above a specified threshold value were subjected to ranking by the cytoHubba plugin (v.3.9.1) (Chin et al., 2014) of Cytoscape (Shannon et al., 2003). As cytoHubba employs 12 different ranking methods, we considered the DEG being top-ranked by ≥6 different methods (i.e., 50% of the total ranking methods) for further investigation. These top-ranked shortlisted proteins were then subjected to local cluster analysis through STRING, and those that were part of multiple clusters were termed potential ‘hubs’ that can be investigated for additional validation of their targetability. The term ‘hub’ refers to a gene or protein that interacts with multiple other genes/proteins. The identified hubs were then subjected to co-occurrence analysis across multiple pathogen genomes to see whether an antibacterial agent targeting these hubs is likely to exert a broad-spectrum activity. This sequence of analysis allowed us to end up with a limited number of proteins that satisfied multiple statistical and biological significance criteria simultaneously: (i) log fold-change ≥2; (ii) FDR≤0.05; (iii) relatively higher node degree; (iv) top-ranking by at least 6 cytoHubba methods; (v) member of more than 1 local network cluster.

### 2.12. Polymerase Chain Reaction (RT-PCR)

Differential expression of the potential hubs identified through network analysis of DEG revealed from WTA was confirmed through PCR too. Primer designing for the selected genes was accomplished through Primer3 Plus (Untergasser et al., 2007). Primer sequences thus obtained were confirmed for their binding within the whole *S. aureus* genome exclusively to the target gene sequence. Primer sequences for all the target genes are listed in Table 1. RNA extraction and quality check was done as described in preceding section. cDNA was obtained by using SuperScript™ VILO™ cDNA Synthesis Kit (Invitrogen Biosciences). PCR assay was conducted by using gene specific primers procured from Sigma-Aldrich. The gene *rpsU* was kept as an endogenous control. The reaction mix used was FastStart Essential DNA Green Master mix (Roche; 06402712001). Real time PCR assay was performed on Quant studio 5 real time PCR machine (Thermo Fisher Scientific, USA). Temperature profile employed is provided in Table S3.

**Table 1.**
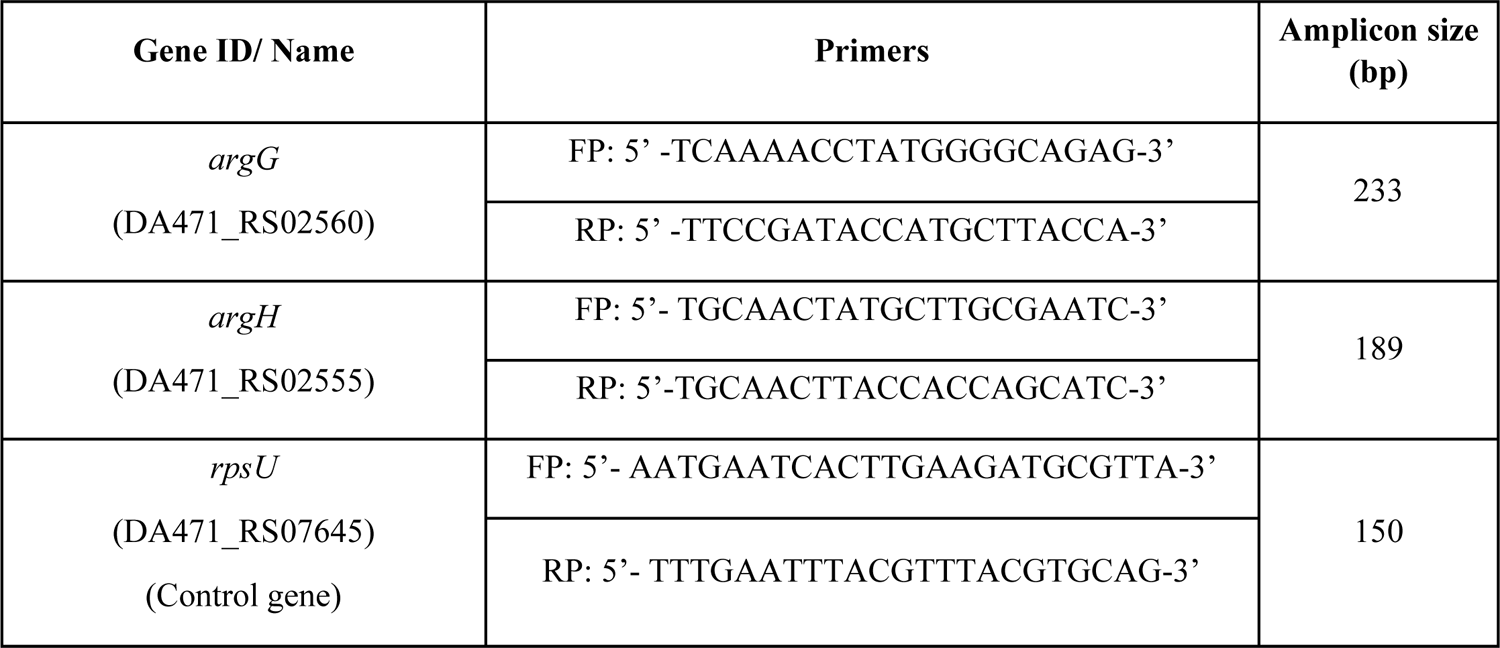
Primer sequences for the target genes.

### 2.13. Statistics

All results reported are means of three or more independent experiments, each performed in triplicate. Statistical significance was assessed through *t*-test performed using Microsoft Excel^®^, and data with *p*≤0.05 was considered to be significant.

## 3. Results and Discussion

### 3.1. Growth inhibitory effect of Silversol against *S. aureus* follows a non-linear dose-response pattern

Silversol exerted its growth inhibitory effect against *S. aureus* at concentrations ≥5 ppm (Figure 2A). Interestingly, the dose-response curve assumed different shapes over different concentration ranges (Figure 2A inset box), considering ‘inhibition of growth’ as ‘response’. For example, in the 1-10 ppm range, it followed a pattern following threshold model, wherein 5 ppm was the minimum concentration required to generate the response. In the 7-15 ppm range it assumed an ‘inverted U’ shape, while in the 10-25 ppm range, it assumed a ‘U’ shape. These different patterns of dose-response relationship can be appreciated better by looking at relevant explanation in Calabrese (2004), Iavicoli et al (2021), etc. The non-linear nature of this dose-response curve stemmed from the fact that some of the higher concentrations (15-20 ppm) were less effective at inhibiting bacterial growth than some of the lower concentrations (8-10 ppm) (Supplementary Image: Figure S4). The minimum concentration of Silversol required to achieve complete visible inhibition of bacterial growth was 10 ppm, and hence it can be considered as MIC. However, growth-inhibitory effect of Silversol till 25 ppm can largely be said to be bacteriostatic, as the cells exposed to these concentrations when plated on fresh media (devoid of Silversol) were able to give rise to lawn growth (Figure S5). Concentrations of Silversol ≥50 ppm exerted bactericidal effect, and the cells exposed to these concentrations when plated on fresh media (devoid of Silversol) could give rise only to few isolated colonies. Non-cidal action of Silversol at lower concentrations was also confirmed in the growth curve experiment, wherein the bacterium was able to partially overcome the growth-inhibitory effect of Silversol, if allowed extended incubation (Figure 2B). However, the maximum cell density achieved by *S. aureus* in presence of 8 ppm and 10 ppm Silversol was 1.71 and 2.03fold lesser than that of control. Onset of growth was much delayed in presence of 10 ppm Silversol. While the control culture was still in exponential phase of growth at 32^nd^ hour of incubation, death phase had already started in the experimental culture (8 ppm Silversol) by 26^th^ hour. Silversol’s 8 ppm level was found to inhibit growth of *S. aureus* nearly by 50%, and hence considering this as ∼IC_50,_ all further experiments (unless specified otherwise) were conducted at this concentration.

**Figure 2.**
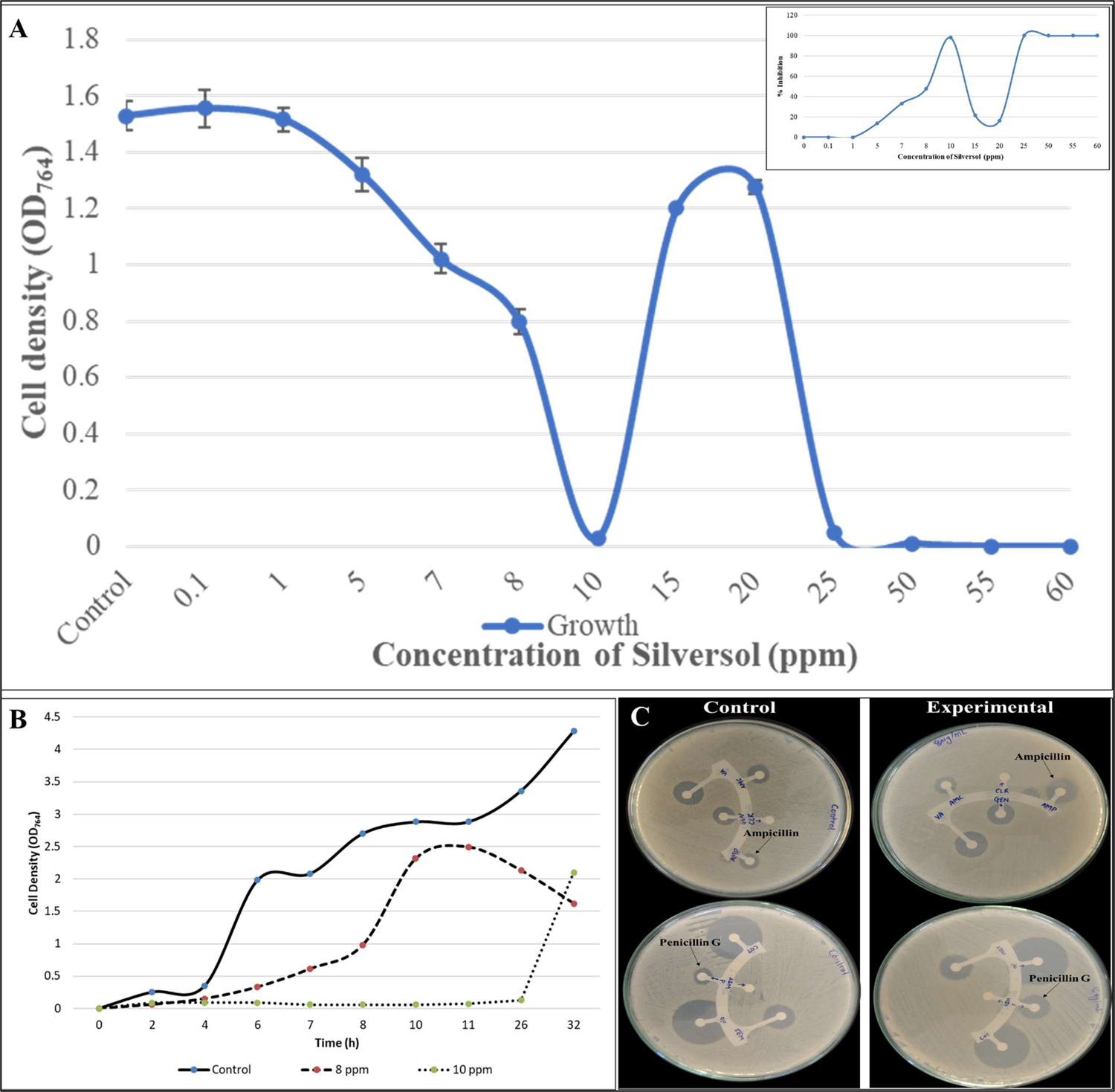
Silversol inhibits growth of *S. aureus,* and enhances its susceptibility towards β-lactams. (A) Silversol inhibits growth of *S. aureus.* Bacterial growth was measured as OD_764_. Vancomycin (5 µg/mL) employed as positive control inhibited bacterial growth by 100%. All the OD points plotted differed from the control at p<0.01. Inset box has plotted percent inhibition vs. time for a better idea of the shape of dose-response curve. (B) Growth curve of *S. aureus* in presence or absence of Silversol. *S. aureus* could achieve lesser growth rate and cell density in presence of bacteriostatic concentrations of Silversol. **(C) Silversol-pre-treatment enhances bacterial susceptibility to beta-lactam antibiotics**. Silversol-pre-treated cells (experimental panel) can be seen surrounded by larger zones of inhibition encircling discs of penicillin and ampicillin.

### 3.2. Silversol pretreatment renders *S. aureus* more susceptible to ampicillin and penicillin

**G.** When Silversol pretreated *S. aureus* cells were subjected to antibiotic susceptibility assay through disc diffusion method, their susceptibility to ampicillin and penicillin was increased by approximately 1.5-fold (Table 2; Figure 2C). To nullify any possible role of ‘post-antibiotic effect’, we did confirm that Silversol-pre-treated cells when transferred into fresh media (devoid of silver), they grow at the same rate as that of control (data not shown). This resistance-modifying activity of Silversol becomes important in light of the fact that strains of *S. aureus* exhibiting beta-lactam resistance have established a considerable ecological niche among human pathogens (Fuda et al., 2005). As approximately only 10% of *S. aureus* clinical isolates (in the United States) are susceptible to penicillin (https://www.cdc.gov/hai/settings/lab/lab_mrsa.html), resistance-modifiers making the bacteria more susceptible to one or more beta-lactams can help widen the utility spectrum of classical antibiotics.

**Table 2.** *Staphylococcus aureus* challenged with antibiotic following pre-treatment with Silversol.

| Antibiotic | Concentration (μg) | Diameter of zone of inhibition (mm) |  | %Difference |
| --- | --- | --- | --- | --- |
|  |  | Control | Experimental |  |
| Penicillin G | 10 | 14±1.14 | 20.5±2.12 | 46.42** |
| Oxacillin | 1 | 23±7.07 | 25.5±6.36 | Non-significant |
| Erythromycin | 15 | R | R |  |
| Clindamycin | 2 | R | R |  |
| Linezolid | 30 | 29.5±3.53 | 30±2.12 |  |
| Co- Trimoxazole | 25 | 35±0 | 35±0 |  |
| Vancomycin | 30 | 20.5±2.12 | 20±0.70 |  |
| Tetracycline | 30 | 29±1.14 | 31±1.41 |  |
| Chloramphenicol | 30 | 28±1.41 | 28±1.41 |  |
| Gentamicin | 10 | 23±12.02 | 23.5±9.19 |  |
| Azithromycin | 15 | R | R |  |
| Ofloxacin | 5 | 35±0 | 36±1.41 |  |
| Methicillin | 5 | 26.5±9.19 | 23.5±4.94 |  |
| Amoxycillin/Clavulanic acid | 30 | 14±0 | 16±2.82 |  |
| Clarithromycin | 15 | R | R |  |
| Ampicillin | 10 | 14±1.41 | 21.5±2.12 | 53.57* |
| Amikacin | 30 | 22.5±2.12 | 23±2.82 | Non-significant |
| Cephalothin | 30 | 30±7.07 | 31±5.65 |  |
| Novobiocin | 5 | 36±1.41 | 36±1.41 |  |
| Teicoplanin | 10 | 21±1.41 | 21.5±0.70 |  |

Antibiotic susceptibility profile of the bacterium was generated using the antibiotic discs- Icosa G-I Plus (HiMedia, Mumbai), through disc diffusion assay performed as per CLSI guidelines. 'R': Resistant i.e. no zone of inhibition was observed. \* $p \leq 0.05$ , \*\* $p \leq 0.01$

### 3.3. Silversol inhibits biofilm formation and renders preformed biofilm non-viable

*S. aureus* in presence of non-growth inhibitory concentration of Silversol could form 59% lesser biofilm than control cells. Neither pre-treatment with Silversol could compromise bacterial ability to form biofilm, nor Silversol could eradicate pre-formed biofilm. However, Silversol brought down the biofilm viability to zero, when added onto pre-formed biofilm (Figure 3A). These results indicate that Silversol could penetrate the biofilm matrix to exert its effect on metabolic activity/ viability of constituent cells. Reduced biofilm formation and decrease in biofilm viability in presence of Silversol was earlier reported by us in case of *Pseudomonas aeruginosa* too (Gajera et al., 2023b). This anti-biofilm activity of Silversol is important considering that bacterial biofilms exert antibiotic tolerance at higher level, and biofilm infections are more difficult to treat (Zhao et al., 2023). Nanoparticles of antibacterial preparations have been demonstrated to possess notable efficacy against bacterial biofilms in a variety of bacterial species (Mishra et al., 2023), and results of the present study corroborate with the same. Anti-biofilm activity of Silversol can be one of the attributes of this formulation making it effective at wound-disinfection, as biofilm-infected wounds take longer to heal (Diban et al., 2023) in absence of an efficient anti-biofilm therapy.

**Figure 3.**
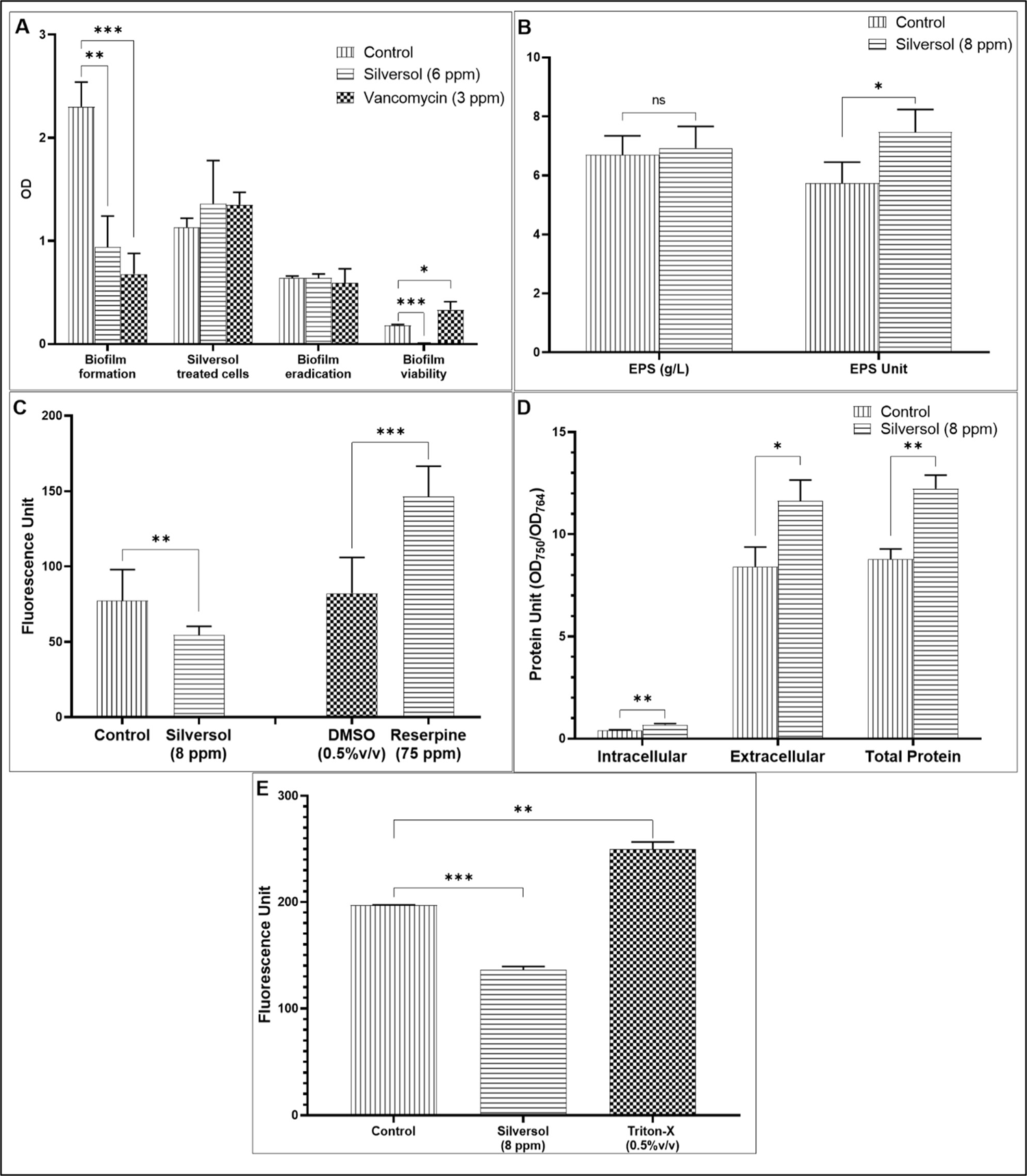
Silversol modulates various important physiological and phenotypic traits in *S. aureus*. (A) Silversol inhibits biofilm formation in *S. aureus* and renders preformed biofilm metabolically inactive. Non-inhibitory concentration of Silversol allowed *S. aureus* to form lesser biofilm (first group of bars); Silversol-pre-treated cells formed biofilm at par to the control cells; When Silversol was added onto pre-formed biofilm, it was not able to eradicate it (third group of bars), but reduced its metabolic activity to non-detectable level (fourth group of bars). Vancomycin employed as a positive control at sub-MIC level inhibited growth by 26%±0.12 and biofilm formation by 70%±12.22. Vancomycin when added onto pre-formed biofilm, could not eradicate biofilm, but increased its metabolic activity by 83%±30.8. Crystal violet assay was performed to quantify biofilm formation, and biofilm viability in the biofilm was estimated through MTT assay. (B) Silversol enhances exopolysaccharide (EPS) synthesis in *S. aureus*. Though there are lesser number of cells in the Silversol-supplemented media, they synthesized EPS in amount equal to their Silversol-unexposed counterparts (first group of bars). However, when EPS Unit was calculated as Cell Density (OD_764_): EPS (g/L) ratio, this was found to be 30%±6 higher in presence of Silversol. (C) Silversol triggers overexpression of efflux machinery in *S. aureus*. Fluorescence of the intracellularly accumulated EtBr was 29%± 5.77 lesser in Silversol-exposed *S. aureus*. Higher (78%±20.12) intracellular accumulation of EtBr was observed in presence of the known efflux inhibitor, reserpine. (D) *S. aureus* grown in presence of Silversol registered higher protein synthesis and secretion. Intracellular and extracellular protein concentrations in *S. aureus* grown in presence of Silversol at sub-MIC level were significantly higher (61±12.7% and 38%±13.9 respectively) as compared to its Silversol-non-exposed counterpart. Protein Unit (i.e., Protein concentration: Cell density ratio) was calculated to nullify any effect of cell density on protein synthesis. (E) Silversol-exposure reduces membrane permeability of *S. aureus*. Silversol-pre-exposed cells allowed 30%±3.18 lesser propidium iodide (PI) to enter them. Cells pre-exposed to Triton-X, a known disruptor of bacterial membranes, allowed 26%±6.97 more PI to enter, as indicated by the fluorescence of intracellularly accumulated dye. *p ≤ 0.05, **p ≤ 0.01, ***p≤0.001

### 3.4. Silversol makes *S. aureus* synthesize and secrete more exopolysaccharide (EPS)

EPS is an important component of biofilm matrix, and it enables the bacterial pathogen for effective adherence to the host and colonize there (Flemming, 2016; Wang et al., 2023). EPS matrix can also protect the pathogen from a variety of challenges including host defense factors (Mielnichuk et al., 2024). Silversol-pre-exposed *S. aureus* culture had 1.30-fold higher EPS production (Figure 3B). Enhanced EPS secretion may be taken as a response to the stress induced in bacterial population by Silversol, as EPS production is known to be a part of bacterial stress-response (Wang et al., 2020; Chen et al., 2022).

### 3.5. Silversol triggers overexpression of efflux machinery in *S. aureus*

*S. aureus* cells accumulated lesser EtBr, when incubated with Silversol than when in Silversol-free media (Figure 3C). This lesser intracellular accumulation of EtBr can be said to have resulted from overexpression of efflux machinery in Silversol-exposed *S. aureus*. While efflux pumps play important role in detoxification, their overexpression may have a negative effect on bacterial physiology (Sun et al., 2014), as overactive efflux machinery will export even some of the essential intracellular content. Results of this *in vitro* efflux assay corroborates well with overexpression of efflux-associated genes (*terC* and *mepA*) in Silversol-exposed *S. aureus* described later.

### 3.6. Protein synthesis and export in *S. aureus* gets enhanced in presence of Silversol

Intracellular and extracellular protein content in *S. aureus* culture grown in presence of Silversol was found to be 1.61-fold and 1.38-fold higher respectively than control (Figure 3D). This indicates that not only overall protein synthesis, but also its export was increased under influence of Silversol. This increased protein export may have arisen from compromised membrane integrity and/or overexpression of efflux machinery (Brown and Skurray, 2001; Pagès et al., 2005; Lennen et al., 2013). It may be said that bacteria respond to the inhibitory effect of such antimicrobials acting as suppressors of protein synthesis, by elevating its protein synthesis and/or secretion machinery to compensate the inhibitory effect of the antibacterial agents (Singh et al., 2001). Such upregulation of protein synthesis can be believed to have stemmed from the translational reprogramming in the stressed cells (Liu and Qian, 2014), as they are dealing with the stress of the antibacterial activity of the Silversol.

### 3.7. Silversol alters membrane permeability of *S. aureus*

Membrane permeability of bacterial cells was quantified in terms of their ability to uptake the fluorescent dye propidium iodide. Lesser uptake of this dye by bacterial cells in presence of Silversol indicates this formulation’s ability to alter membrane permeability (Figure 3E). Reduced permeability might have compromised bacterial capacity to allow intake of nutrients leading to stunted growth. This combined with overexpression of efflux function (Figure 3C) and protein secretion (Figure 3D) can be expected to force bacteria to face scarcity of essential nutrients and metabolites. Alterations in membrane functioning can affect multiple bacterial traits such as surface charge, permeability, fluidity, and stability of the bacterial membrane, antibiotic susceptibility etc. (Nikolic and Mudgil, 2023). A correlation between antibacterial activity of Silversol and its ability to alter membrane fluidity can not be ruled out, as expected of other membranotropic agents (Olchowik-Grabarek et al., 2022).

### 3.8. Silversol causes multiple genes in *S. aureus* to express differentially

After confirming the growth-inhibitory activity of Silversol against *S. aureus*, we compared the gene expression profile of the Silversol-exposed *S. aureus* with that of control at the whole transcriptome level. Whole transcriptome analysis identified a total of 90 differentially expressed genes (3.32% of total genome) in Silversol-treated *S. aureus*, of which 49 were down-regulated (Table 3) and 41 were up-regulated (Table 4). Corresponding heat map (Figure S6) and volcano plot (Figure S7) can be seen in supplementary file. A summary of function-wise categorization of all the differentially expressed genes (DEG) is presented in Figure 4. Among the down-regulated DEG, the one with highest log fold change (logFC 5.14) value was *CidR,* which is the transcriptional regulator of the LysR family. This regulator is known to control stationary phase cell death in *S. aureus* (Chaudhari et al., 2016) by influencing the expression of operons that display pro- and anti-death functions. Since primary role of CidR regulon is to limit acetate-dependent potentiation of cell death in staphylococcal populations, its down-regulation can be expected to have negative effect on bacterial growth. The *CidR* gene besides regulating stationary-phase survival of *S. aureus*, also affects antibiotic tolerance in this bacterium (Yang et al., 2005).

**Figure 4.**
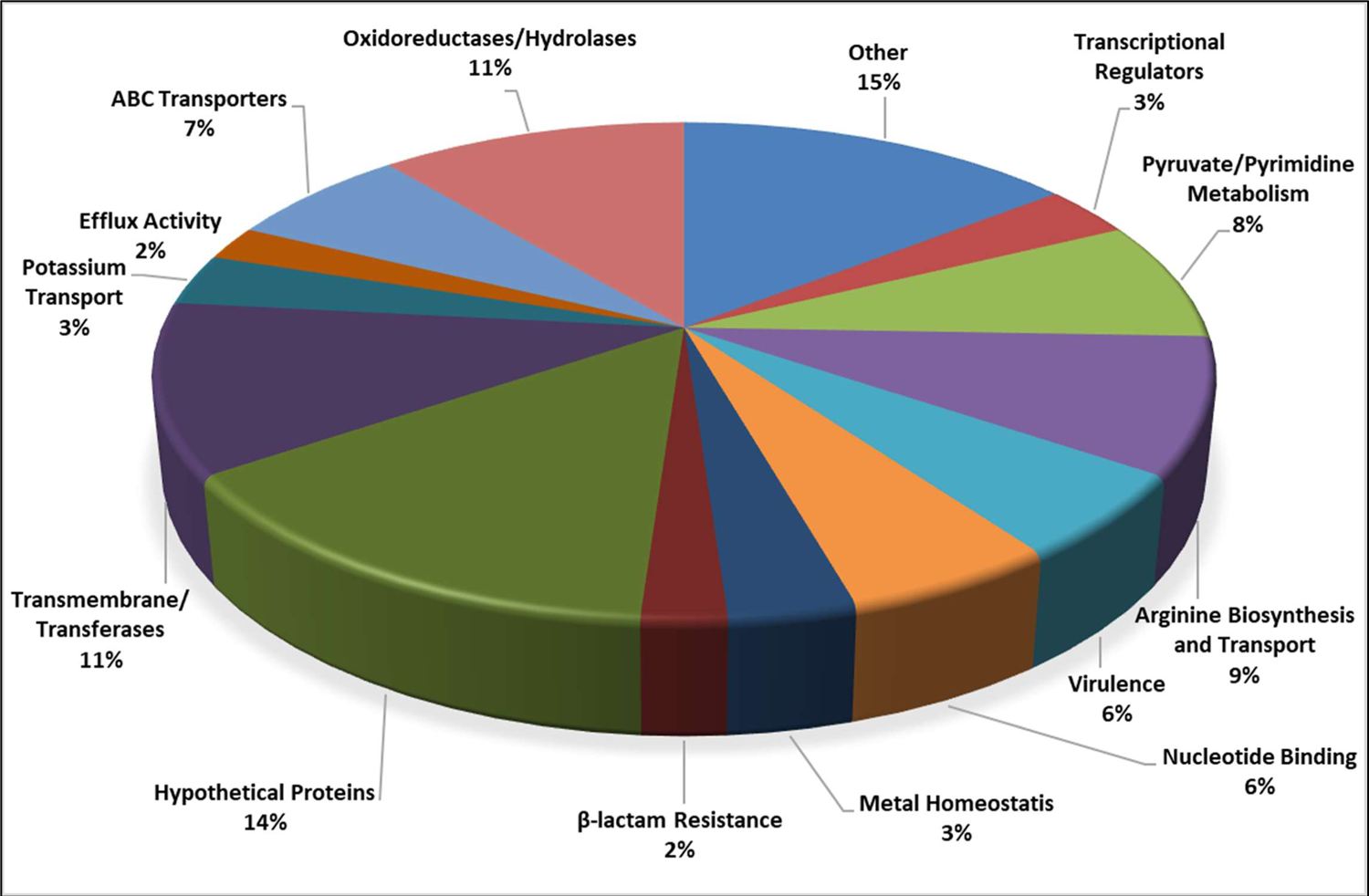
Function-wise categorization of the significantly differentially expressed genes in Silversol-treated *Staphylococcus aureus*(% values represent the fraction of the total number of DEG)

**Table 3.**
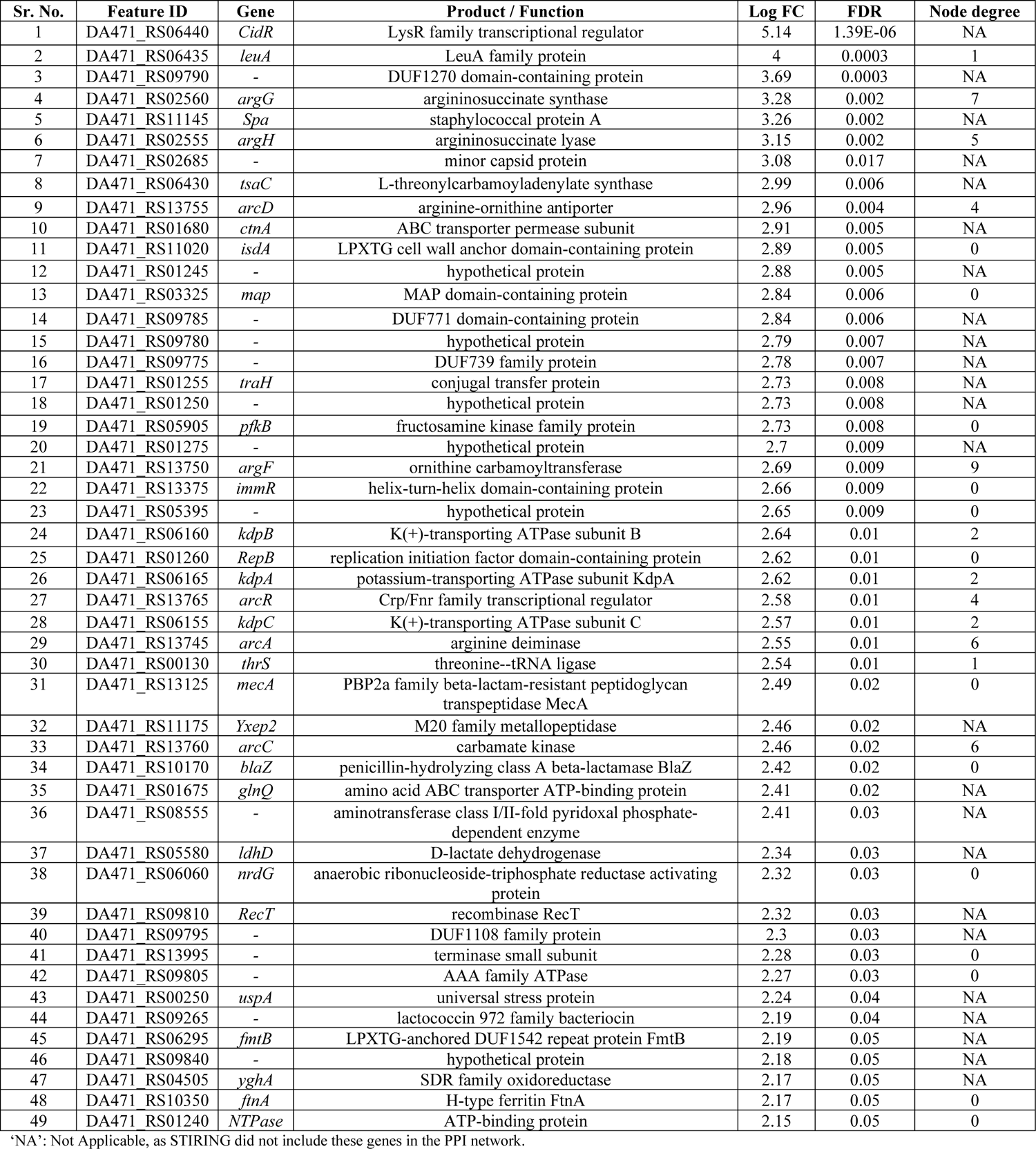
List of down-regulated genes in Silversol exposed *S. aureus* satisfying the dual criteria of log fold change ≥2 and FDR≤0.05.

| Sr. No. | Feature ID | Gene | Product / Function | Log FC | FDR | Node degree |
| --- | --- | --- | --- | --- | --- | --- |
| 1 | DA471_RS06440 | *CidR* | LysR family transcriptional regulator | 5.14 | 1.39E-06 | NA |
| 2 | DA471_RS06435 | *leuA* | LeuA family protein | 4 | 0.0003 | 1 |
| 3 | DA471_RS09790 | - | DUF1270 domain-containing protein | 3.69 | 0.0003 | NA |
| 4 | DA471_RS02560 | *argG* | argininosuccinate synthase | 3.28 | 0.002 | 7 |
| 5 | DA471_RS11145 | *Spa* | staphylococcal protein A | 3.26 | 0.002 | NA |
| 6 | DA471_RS02555 | *argH* | argininosuccinate lyase | 3.15 | 0.002 | 5 |
| 7 | DA471_RS02685 | - | minor capsid protein | 3.08 | 0.017 | NA |
| 8 | DA471_RS06430 | *tsaC* | L-threonylcarbamoyladenylate synthase | 2.99 | 0.006 | NA |
| 9 | DA471_RS13755 | *arcD* | arginine-ornithine antiporter | 2.96 | 0.004 | 4 |
| 10 | DA471_RS01680 | *ctnA* | ABC transporter permease subunit | 2.91 | 0.005 | NA |
| 11 | DA471_RS11020 | *isdA* | LPXTG cell wall anchor domain-containing protein | 2.89 | 0.005 | 0 |
| 12 | DA471_RS01245 | - | hypothetical protein | 2.88 | 0.005 | NA |
| 13 | DA471_RS03325 | *map* | MAP domain-containing protein | 2.84 | 0.006 | 0 |
| 14 | DA471_RS09785 | - | DUF771 domain-containing protein | 2.84 | 0.006 | NA |
| 15 | DA471_RS09780 | - | hypothetical protein | 2.79 | 0.007 | NA |
| 16 | DA471_RS09775 | - | DUF739 family protein | 2.78 | 0.007 | NA |
| 17 | DA471_RS01255 | *traH* | conjugal transfer protein | 2.73 | 0.008 | NA |
| 18 | DA471_RS01250 | - | hypothetical protein | 2.73 | 0.008 | NA |
| 19 | DA471_RS05905 | *pfkB* | fructosamine kinase family protein | 2.73 | 0.008 | 0 |
| 20 | DA471_RS01275 | - | hypothetical protein | 2.7 | 0.009 | NA |
| 21 | DA471_RS13750 | *argF* | ornithine carbamoyltransferase | 2.69 | 0.009 | 9 |
| 22 | DA471_RS13375 | *immR* | helix-turn-helix domain-containing protein | 2.66 | 0.009 | 0 |
| 23 | DA471_RS05395 | - | hypothetical protein | 2.65 | 0.009 | 0 |
| 24 | DA471_RS06160 | *kdpB* | K(+)-transporting ATPase subunit B | 2.64 | 0.01 | 2 |
| 25 | DA471_RS01260 | *RepB* | replication initiation factor domain-containing protein | 2.62 | 0.01 | 0 |
| 26 | DA471_RS06165 | *kdpA* | potassium-transporting ATPase subunit KdpA | 2.62 | 0.01 | 2 |
| 27 | DA471_RS13765 | *arcR* | Crp/Fnr family transcriptional regulator | 2.58 | 0.01 | 4 |
| 28 | DA471_RS06155 | *kdpC* | K(+)-transporting ATPase subunit C | 2.57 | 0.01 | 2 |
| 29 | DA471_RS13745 | *arcA* | arginine deiminase | 2.55 | 0.01 | 6 |
| 30 | DA471_RS00130 | *thrS* | threonine--tRNA ligase | 2.54 | 0.01 | 1 |
| 31 | DA471_RS13125 | *mecA* | PBP2a family beta-lactam-resistant peptidoglycan transpeptidase MecA | 2.49 | 0.02 | 0 |
| 32 | DA471_RS11175 | *Yxep2* | M20 family metallopeptidase | 2.46 | 0.02 | NA |
| 33 | DA471_RS13760 | *arcC* | carbamate kinase | 2.46 | 0.02 | 6 |
| 34 | DA471_RS10170 | *blaZ* | penicillin-hydrolyzing class A beta-lactamase BlaZ | 2.42 | 0.02 | 0 |
| 35 | DA471_RS01675 | *glnQ* | amino acid ABC transporter ATP-binding protein | 2.41 | 0.02 | NA |
| 36 | DA471_RS08555 | - | aminotransferase class I/II-fold pyridoxal phosphate-dependent enzyme | 2.41 | 0.03 | NA |
| 37 | DA471_RS05580 | *ldhD* | D-lactate dehydrogenase | 2.34 | 0.03 | NA |
| 38 | DA471_RS06060 | *nrdG* | anaerobic ribonucleoside-triphosphate reductase activating protein | 2.32 | 0.03 | 0 |
| 39 | DA471_RS09810 | *RecT* | recombinase RecT | 2.32 | 0.03 | NA |
| 40 | DA471_RS09795 | - | DUF1108 family protein | 2.3 | 0.03 | NA |
| 41 | DA471_RS13995 | - | terminase small subunit | 2.28 | 0.03 | 0 |
| 42 | DA471_RS09805 | - | AAA family ATPase | 2.27 | 0.03 | 0 |
| 43 | DA471_RS00250 | *uspA* | universal stress protein | 2.24 | 0.04 | NA |
| 44 | DA471_RS09265 | - | lactococcin 972 family bacteriocin | 2.19 | 0.04 | NA |
| 45 | DA471_RS06295 | *fmtB* | LPXTG-anchored DUF1542 repeat protein FmtB | 2.19 | 0.05 | NA |
| 46 | DA471_RS09840 | - | hypothetical protein | 2.18 | 0.05 | NA |
| 47 | DA471_RS04505 | *yghA* | SDR family oxidoreductase | 2.17 | 0.05 | NA |
| 48 | DA471_RS10350 | *ftnA* | H-type ferritin FtnA | 2.17 | 0.05 | 0 |
| 49 | DA471_RS01240 | *NTPase* | ATP-binding protein | 2.15 | 0.05 | 0 |

‘NA’: Not Applicable, as STIRING did not include these genes in the PPI network.

**Table 4.**
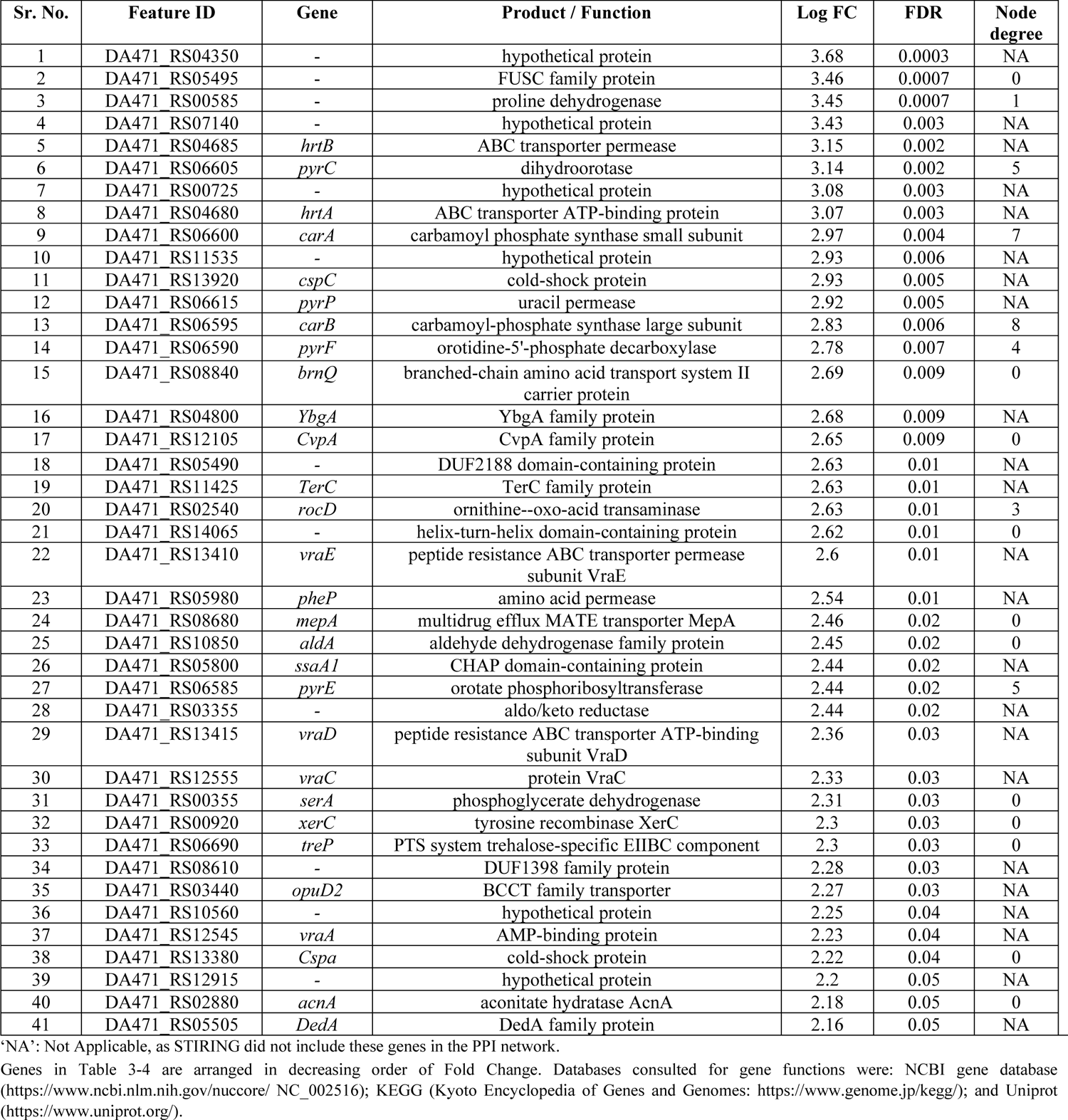
List of up-regulated genes in Silversol exposed *S. aureus* satisfying the dual criteria of log fold change ≥2 and FDR≤0.05.

Another heavily down-regulated gene in the Silversol-exposed *S. aureus* was *spa* (3.26↓) coding for staphylococcal protein A. Latter is a multifunctional virulence factor of *S. aureus*, which plays role in the inhibition of the host innate and adaptive immune responses. Owing to its immunoglobulin-binding capacity, this protein confers protection on *S. aureus* from phagocytic killing via inhibition of Ig Fc region. Additionally, spa is known to prevent the host elicited B-cell response, decreasing long-term antibody production, inhibit osteogenesis, and act as a pro-inflammatory factor in the lung (Rigi et al., 2019). Protein A has also been identified as an essential component of the *S. aureus* biofilm, which induces bacterial aggregation in liquid medium and during biofilm formation. Besides interacting with multiple immunologically important eukaryotic receptors, protein A contributes considerably toward the development of biofilm-associated infections (Merino et al., 2009), and hence its downregulation by Silversol can be of high clinical relevance.

In addition to spa, other virulence associated genes downregulated in Silversol-exposed *S. aureus* were *TraH* (2.73↓), and the one coding for ABC transporter permease subunit (2.91↓). Latter is a part of the ABC transporter complex CntABCDF involved in the uptake of metal, particularly in the import of divalent metal ions like nickel, cobalt and zinc. Its downregulation may be expected to disturb metal homeostasis in the bacterial populations as many important enzymes rely on their metal cofactors for proper functioning. This permease also is believed to contribute towards virulence of *S. aureus*, as it has been shown to be necessary for full urease activity *in vitro* (Remy et al., 2013), and urease is well recognized as a virulence factor in multiple pathogenic bacteria (Rutherford, 2014; Aliyeva-Schnorr et al., 2023; Oliyaei et al., 2023).

TraH mentioned in preceding paragraph is a conjugal transfer protein, and a component of the Type IV secretion system (T4SS). Since for many gram-positive pathogens, conjugative plasmid transfer is an important means of spreading antibiotic resistance, components of T4SS are being viewed as potential targets for alternative anti-pathogenic strategies (Laverde et al., 2017).

Among other important gene clusters downregulated in Silversol-exposed *S. aureus*, one notable was the kdp system, which is a potassium uptake system. Three components [kdpA (2.62↓), kdpB (2.64↓), and kdpC (2.57↓)] of this system were found to be significantly downregulated in our experimental culture. *kdpB* has been shown as one of the core genes regulating cell death in *S. aureus* (Yee et al., 2019). Potassium has many key functions within bacterial cells. For example, it is required for the activity of multiple intracellular enzymes, acts as an intracellular second messenger, and is involved in the maintenance of a constant internal pH and membrane potential. One of the clinically important character of *S. aureus* is its ability to survive in/on high-salt areas of human nares and skin is also helped by its capacity of efficient potassium uptake and intracellular accumulation of this cation (Gründling, 2013). Since the Kdp system is widely distributed among bacteria (Xue et al., 2011), any formulation targeting this system can be expected to exert a broad-spectrum antibacterial activity, and in fact silver nanoparticles have been demonstrated to possess a broad activity spectrum in terms of their ability to inhibit growth of different gram-positive and gram-negative bacteria (Guzman et al., 2012; Dominguez et al., 2020).

Downregulation of two of the genes, *mecA* and *blaZ*, associated with beta-lactam resistance corroborated well with results of antibiotic susceptibility (Figure 2C and Table 2), wherein Silversol-preexposure was found to increase the susceptibility of *S. aureus* to beta-lactams. Occurrence of *mecA* and *blaZ* genes has been reported in methicillin-resistant *S. aureus* associated with vaginitis among pregnant women (Okiki et al., 2020). Expression of these genes may be considered important for expression of antibiotic-resistant phenotype in clinical isolates, as functionality of at least one *mecA* regulator is necessary for *S. aureus* to be oxacillin-resistant (Rosato et al., 2003). Penicillin resistance in *S. aureus* is manifested predominantly via the production of β-lactamase encoded by the *blaZ* gene, and testing for presence of this gene is also recommended by the Clinical and Laboratory Standards Institute for cases of serious *S. aureus* infection (Pereira et al., 2014).

Among the top ten up-regulated genes in Silversol-exposed *S. aureus*, two were *hrtA* (3.07↑) and *hrtB* (3.15↑). Four more genes (*vraA, vraC, vraD*, and *vraE*) coding for components of the ABC transporter complex hrt were also upregulated. HrtAB system is a hemin-regulated ABC transporter that protect *S. aureus* against hemin toxicity. Since *S. aureus* has been reported to launch an altered gene expression program involving differential expression of Hrt genes, when it senses membrane damage; differential upregulation of *hrtA* and *hrtB* in our experimental culture can be taken as an indication of Silversol’s ability to alter membrane permability in this bacterium. Overexpression of *hrtB* can lead to dysregulated pore formation, and its upregulation has also been reported in *S. aureus* challenged with channel-forming antimicrobial peptides (Attia et al., 2010). Altered abundance of Hrt proteins has also been reported in *S. aureus* cultures facing alterations in iron status (Friedman et al., 2006), and hence upregulation of these genes in presence of Silversol can be considered an indication of this formulation’s ability to disturb iron-homeostasis. Silversol’s potential ability to trigger iron-starvation in *S. aureus* can be one of the reasons underlying its successful clinical applications (e.g., wound disinfection), as the role of bacterial iron acquisition during pathogenesis is well established (Sheldon et al., 2016; Zughaier and Cornelis, 2018).

Among the top-25 upregulated genes in Silversol-exposed *S. aureus*, two [*terC* (2.63↑) and *mepA* (2.46↑)] were associated with efflux activity. Efflux machinery besides throwing out toxic items, also has important physiological functions in bacteria (Sun et al., 2014). In general, upregulation of efflux machinery can be taken as a sign of the bacteria making efforts for detoxification, for example in case of this study they may be trying to efflux silver. However, overexpression of efflux function can lead to leakage of even essential molecules from within the cell and thereby affecting bacterial growth and fitness negatively. The way efflux inhibitors are considered as potential anti-pathogenic agents (Ahmad et al., 2018), efflux agonists triggering overexpression of the efflux pumps can also be pursued as a novel antibacterial strategy. Of the two up-regulated genes mentioned above, TerC belongs to a family of integral membrane proteins, and has been implicated in resistance to metal ions like tellurium (Paruthiyil et al., 2020) and Mn (Chen et al., 2024). TerC helps bacteria alleviate metal toxicity by participating in efflux of metal ions. In case of the present study, TerC might have got activated in response to silver overload. Owing to the widespread occurrence of the TerC family proteins among bacteria and its possible influence on host-pathogen interactions, it can be an important antibacterial target (Turkovicova et al., 2016; He et al., 2023). Another efflux-associated upregulated gene *mepA* is a multidrug resistance efflux protein involved in transporting several clinically relevant monovalent and divalent biocides, the fluoroquinolone antibiotics-norfloxacin and ciprofloxacin, and also tigecycline (Poole, 2005). Since mepA has a broad substrate profile (Kaatz et al., 2005), its poorly regulated overexpression (as observed in Silversol-exposed *S. aureus*) can trigger efflux of multiple substrates probably including some essential metabolites. Though metal nanoparticles have been postulated to exert inhibitory action against multidrug resistance efflux pumps (Hasani et al., 2019), their ability to trigger uncontrolled overexpression also needs to be investigated for possible clinical exploitation.

### 3.9. Network analysis of DEG

To have a deeper insight into the interactions among DEG identified in Silversol-exposed *S. aureus*, we subjected all the DEG to network analysis through STRING. The resulting protein-protein interaction (PPI) (Figure 5A) network had 41 nodes connected through 41 edges, with an average node degree of 2. Since the number of edges (41) in this PPI network is almost 3.4-fold higher than expected (12), this network can be said to possess significantly more interactions among the member proteins than what can be expected for a random set of proteins having identical sample size and degree distribution. Such an enrichment is suggestive of the member proteins being at least partially biologically connected. When we arranged all the 41 nodes in decreasing order of node degree, 22 nodes were found to have a non-zero score, and we selected top 13 genes (Table 3-4) with a node degree ≥3 for further ranking by different cytoHubba methods (Table 5). Then we looked for genes which appeared among top ranked candidates by ≥6 cytoHubba methods. This allowed us to shortlist six genes which were ranked among top-6 by ≥9 cytoHubba methods to be taken for further cluster analysis. Interaction map of these 6 important genes (Figure 5B) showed them to be networked with the average node degree score of 4.33. Number of edges possessed by this network was 13 as against expected 1 for any such random set of proteins. These 6 genes were found to be distributed among three different local network clusters. Two of the genes, *argG* and *argH* were part of two of these three clusters, hence we chose them for further RT-PCR validation. PCR assay did confirm differential expression of these two genes in Silversol-exposed *S. aureus* (Figure 6A). Hence arginine metabolism can be believed to be one of the major targets of Silversol in *S. aureus*.

**Figure 5.**
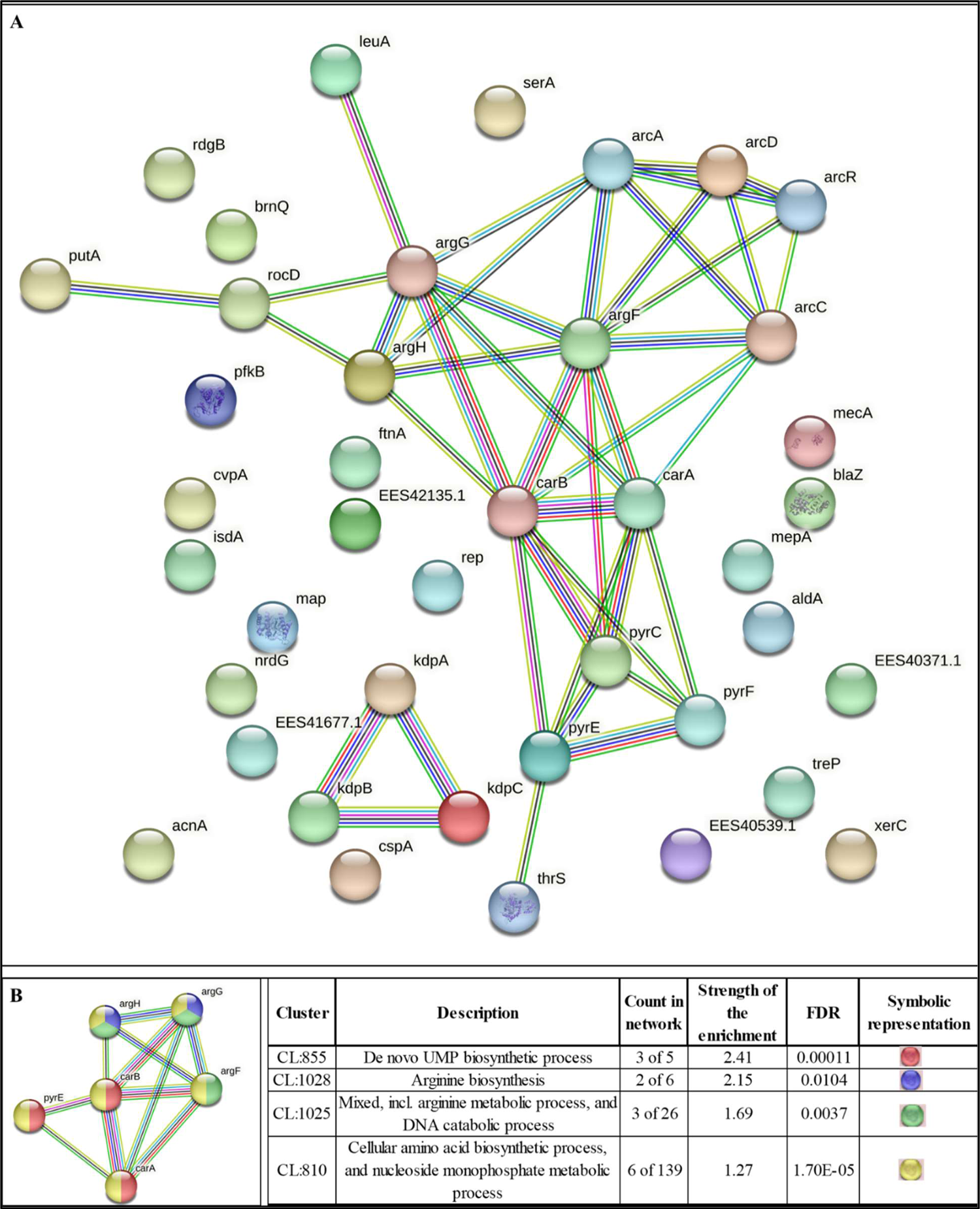
(A) Protein-Protein Interaction (PPI) network of DEG following the dual criteria of fold change ≥ log 2 and FDR ≤ 0.05 in Silversol-exposed *S. aureus.* PPI enrichment *p*–value 3.81e-11. (B) PPI network of top-ranked genes revealed through cytoHubba among DEG in Silversol-exposed *S. aureus.* PPI enrichment *p*–value 1.81e-14.

**Figure 6.**
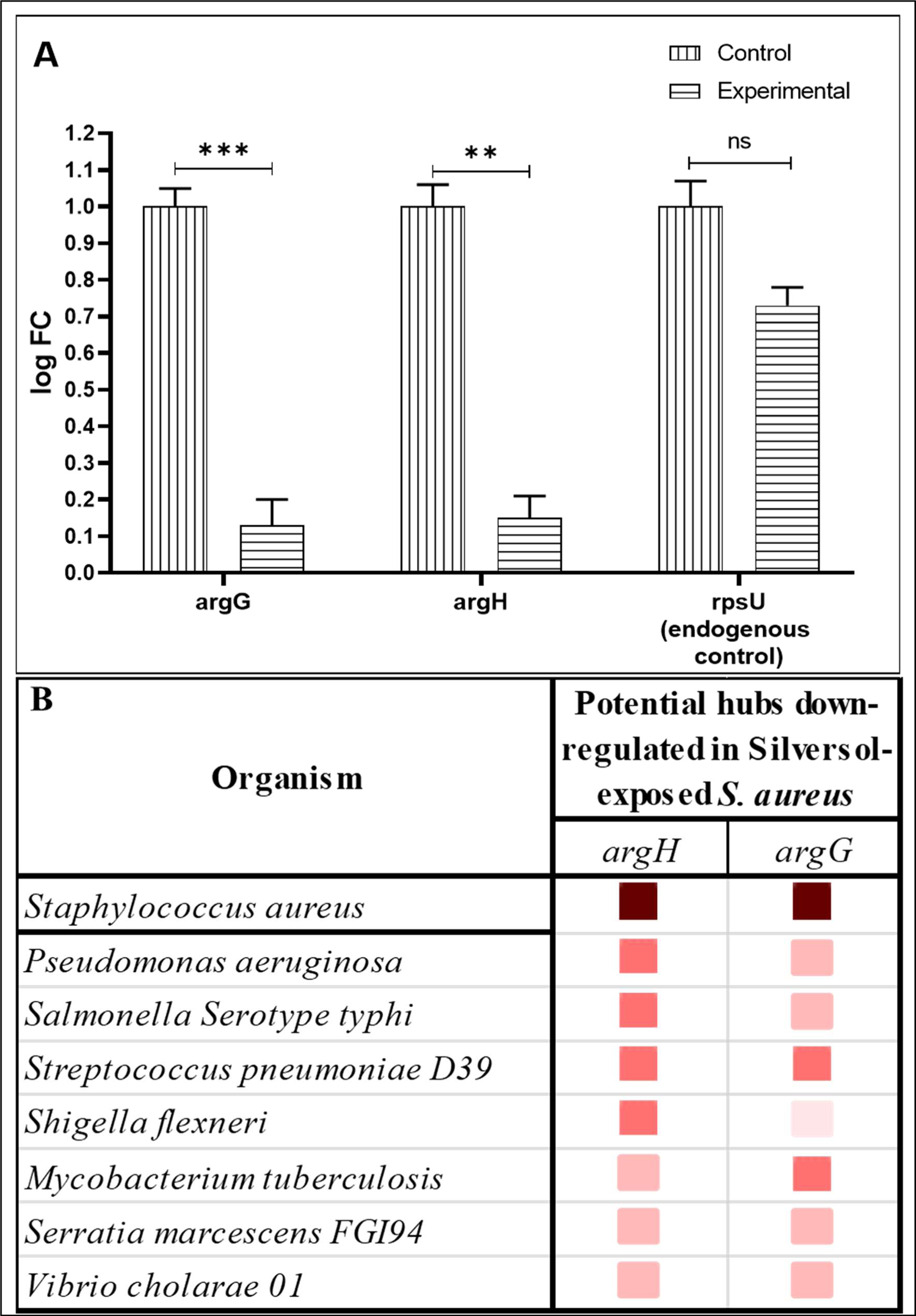
(A) Confirmation of differential expression of selected genes in Silversol-treated *S. aureus* through RT-PCR. (B) Cooccurrence analysis of genes coding for potential targets in *S. **aureus*****, across multiple pathogens**. The darker the shade of the squares, higher is the homology between the genes being compared.

**Table 5.** cytoHubba ranking of the top twelve high node degree genes.

| No. | Gene ID | Gene Name | Number of methods ranking this protein among top 12 | Names of 12 ranking methods of CytoHubba and rank score provided by them |  |  |  |  |  |  |  |  |  |  |  |
| --- | --- | --- | --- | --- | --- | --- | --- | --- | --- | --- | --- | --- | --- | --- | --- |
|  |  |  |  | Degree | MNC | DMNC | MCC | Bottleneck | EcCentricity | Closeness | Radiality | Betweenness | Stress | CC | EPC |
| 1 | 1200 | *carA* | 11 | 8 | 8 | 0.466 | 66 | 1 | 0.5 | 9.5 | 3.090 | 9.810 | 38 | 0.57 | 8.39 |
| 2 | 0956 | *argG* | 10 | 6 | 6 | 0.475 | 36 | 1 | 0.5 | 8.5 | 2.909 | 4.335 | 22 | 0.67 | 7.79 |
| 3 | 0955 | *argH* | 10 | 7 | 7 | 0.439 | 42 | 1 | 0.5 | 9 | 3 | 9.933 | 34 | 0.57 | 8.21 |
| 4 | 1201 | *carB* | 10 | 9 | 9 | 0.405 | 68 | 2 | 0.5 | 10 | 3.181 | 20.516 | 60 | 0.47 | 8.49 |
| 5 | 1165 | *argF* | 9 | 8 | 8 | 0.437 | 50 | 5 | 0.5 | 9.5 | 3.090 | 13.885 | 44 | 0.53 | 8.48 |
| 6 | 1203 | *pyrE* | 9 | 5 | 5 | 0.518 | 30 | 1 | 0.5 | 8 | 2.818 | 1.598 | 8 | 0.8 | 7.39 |
| 7 | 1199 | *pyrC* | 3 | 5 | 5 | 0.518 | 30 | 1 | 0.5 | 8 | 2.818 | 1.598 | 8 | 0.8 | 7.40 |
| 8 | 2906 | *arcA* | 2 | 5 | 5 | 0.324 | 10 | 1 | 0.33 | 7.833 | 2.727 | 4.523 | 16 | 0.5 | 7.45 |
| 9 | 2903 | *arcC2* | 2 | 5 | 5 | 0.388 | 12 | 1 | 0.5 | 8 | 2.818 | 3.216 | 14 | 0.8 | 7.63 |
| 10 | 2902 | *arcR* | 2 | 3 | 3 | 0.308 | 4 | 1 | 0.33 | 6.833 | 2.545 | 0.581 | 4 | 0.67 | 5.93 |
| 11 | 1202 | *pyrF* | 2 | 4 | 4 | 0.568 | 24 | 1 | 0.33 | 7.166 | 2.545 | 0 | 0 | 1 | 6.88 |
| 12 | 0170 | *rocD* | 2 | 3 | 3 | 0.463 | 6 | 1 | 0.33 | 6.666 | 2.454 | 0 | 0 | 1 | 6.32 |

One of the high node degree genes *arcD* was not considered by cytoHubba.

All the six identified hubs were shown by cluster analysis to belong to amino acid biosynthesis (Figure 5B). All these six genes are participants of the arginine biosynthesis pathway. Three of them (*argG, argH*, and *argF*) are involved in urea cycle, which is a part of arginine biosynthesis pathway. Hence amino acid metabolism in general, and arginine metabolism and urea cycle in particular seems to be targeted by Silversol in *S. aureus*.

To have an increased confidence in our finding that one of the major modes of action of Silversol against *S. aureus* is to interfere with arginine synthesis, we did an additional experiment wherein *S. aureus* was incubated either in Silversol- or L-arginine (HiMedia)-supplemented media, as well as in a media containing both Silversol and arginine together. Arginine supplementation allowed bacteria to overcome inhibitory effect of Silversol (Figure 7), confirming Silversol’s ability to interfere with arginine metabolism. This observation becomes relevant in the context that amino acid pathways have recently been targeted as a novel approach to manage bacterial infections. In particular, arginine metabolism has been illustrated to be important for bacterial pathogenesis (Xiong et al., 2016). Since arginine and its metabolites are utilized as energy sources by various pathogens, its reduced availability can trigger nutrient stress. Reduced availability of arginine may also lead to reduced expression of various pathogenicity genes. The importance of arginine metabolism has been reported in multiple pathogens like *Salmonella typhimurium, Helicobacter pylori, Streptococcus pneumoniae*, *Pseudomonas aeruginosa*, and *Mycobacterium tuberculosis* as a source of energy and as a trigger for polyamine synthesis required for efficient pathogenesis. Owing to the critical role of arginine in establishing pathogenesis, several pathogens employ an array of mechanisms for finetuning arginine metabolism (Gogoi et al., 2016), and breach of this finetuning can be expected to compromise bacterial fitness. While pathogens employ strategies to counteract immune responses by interfering with host arginine metabolism, our study shows Silversol to do the same against *S. aureus*. However, Silversol’s growth-inhibitory action against other pathogens may or may not stem from same mode of action. For example, in our recent study (Gajera et al., 2023b) describing Silversol’s anti-*P. aeruginosa* activity, arginine metabolism was not found to be among the physiological traits attacked by this colloidal silver formulation. It is possible, that despite being a broad-spectrum antibacterial agent, mode of action of Silversol against different pathogenic bacteria may differ more or less.

**Figure 7.**
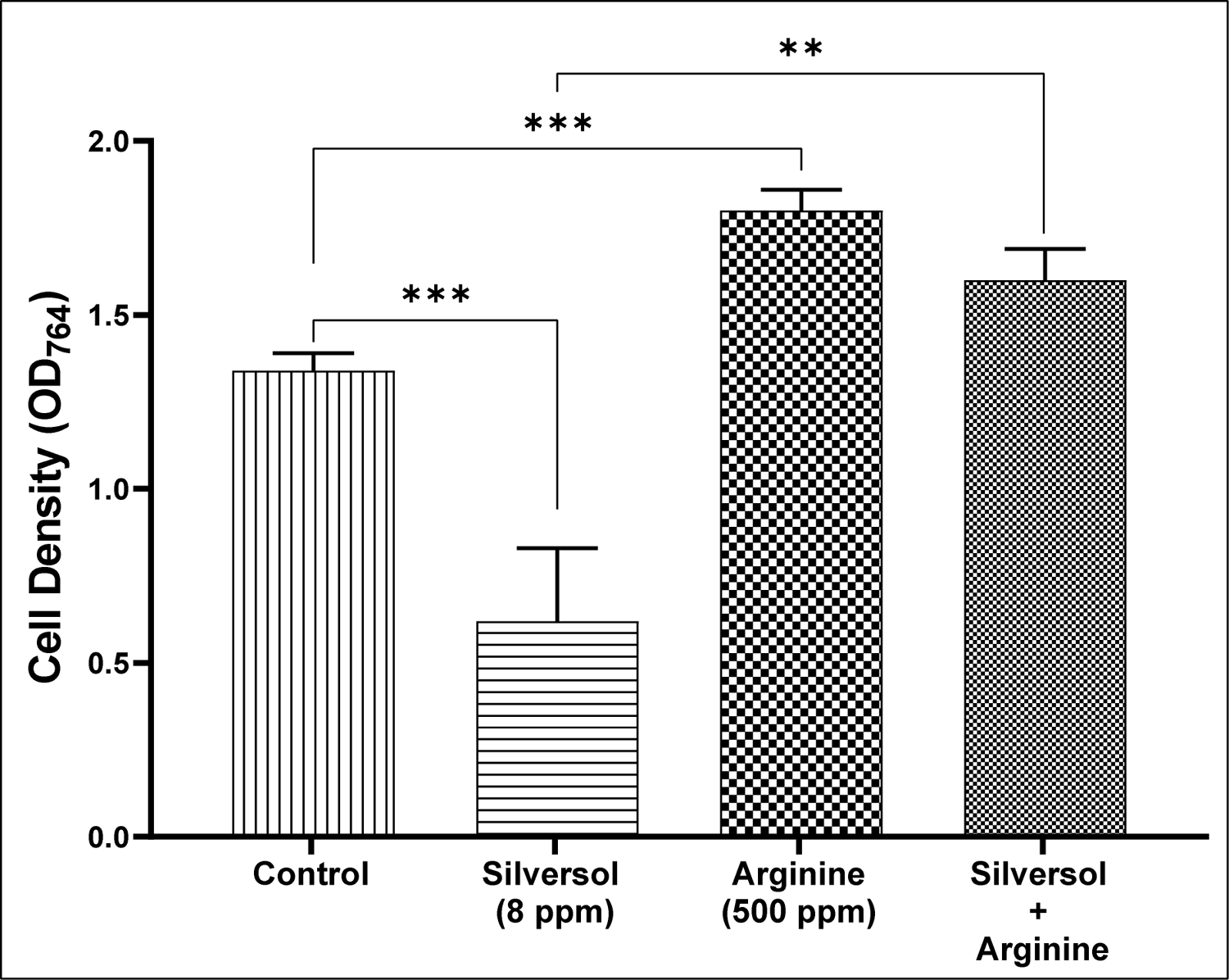
*S. aureus* can resist Silversol’s growth-inhbitory effect in presence of arginine. When *S. aureus* was challenged with Silversol (sub-MIC) in absence or presence of arginine, in latter case Silversol could not show its antibacterial effect. However this protection against Silversol conferred on bacteria by arginine-supplementation of media disappeared when higher concentration of Silversol was used to challenge *S. aureus* (data not shown). It is possible that at higher concentrations bactericidal effect of Silversol might be stemming from additional modes of action, rather than inhibiting arginine biosynthesis.

**Figure 7.**
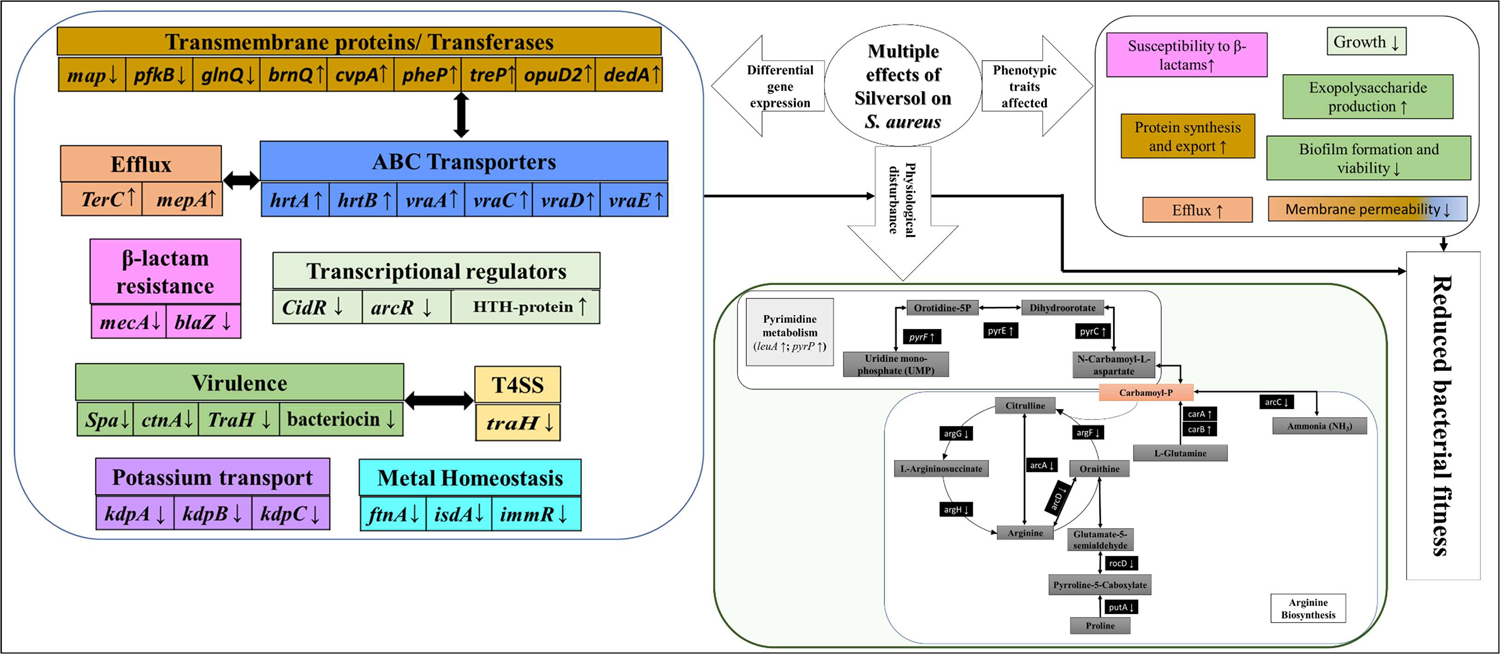
Overall schematic of multiple effects of Silversol on *S. aureus*. Various physiological and phenotypic traits of *S. aureus* affected under the influence of sub-lethal concentration of Silversol are depicted. Boxes with identical colours have mutual functional relevance. (↑) or (↓) arrow indicates up- or down-regulation of the concerned gene.

Arginine has an important role as a substituent for potassium. Under conditions of potassium-limitation, bacteria respond by overproduction of arginine, which may partially substitute for potassium to buffer the negative charge of DNA (Gundlach et al., 2018). However, in presence of Silversol, *S. aureus* seems to have failed to do so, as in our study, the Silversol-exposed *S. aureus* is simultaneously suffering from potassium-limitation as well as downregulation of arginine biosynthesis.

We also conducted a gene cooccurrence pattern analysis of gene families across genomes (through STRING) with respect to the two potential hubs identified by us and confirmed through PCR (Figure 6A). These two genes appeared to have homologues in few other important pathogens too (Figure 6B), particularly *Streptococcus pneumoniae*. Whether Silversol’s antibacterial action against other pathogenic bacteria also involves targeting arginine biosynthesis remains an interesting question to be pursued.

## 4. Conclusion

This study investigated antibacterial effect of a colloidal nanosilver formulation Silversol against antibiotic-resistant *S. aureus*. This formulation was found to inhibit growth of *S. aureus* by targeting a wide variety of genes and physiological traits including efflux, biofilm and exopolysaccharide formation, antibiotic susceptibility, arginine biosynthesis, protein synthesis, potassium uptake, transcriptional regulators, etc. A summary of multiple modes of action of Silversol against *S. aureus* is presented in Figure-8. Particularly arginine metabolism appeared to be one of the major targets of Silversol in *S. aureus*. While aromatic amino acid biosynthesis has been shown as a viable target in *S. aureus* (O’Connell et al., 1993), and *argJ* was shown to be a potential core regulator for *S. aureus* persistence in various stresses (Yee et al., 2019), to the best of our awareness, this is the first report showing *argG* and *argH* as important antibacterial targets in this pathogen. Though antibacterial mechanisms of silver and its nano-formulations have been investigated extensively over past decades, arginine biosynthesis was hitherto not shown to be targeted by silver in susceptible bacteria. Greater insights into arginine metabolism of pathogenic bacteria, and relationship of arginine metabolism with bacterial pathogenesis, would provide possible targets for controlling bacterial infections. Arginine-tagged drug delivery systems can be designed to target specific subcellular locations like pathogen-containing vacuoles inside host cells. In view of the rampant antibiotic resistance, targeting arginine biosynthesis in bacterial pathogens may be a potent approach towards development of next generation anti-pathogenic formulations. Important targets in *S. aureus* attacked by Silversol identified in this study can prove vital input for other antibacterial drug discovery programmes. Amino acid biosynthesis pathways are critical for bacterial growth in nutrient-limiting conditions in the host. Surprising connections between bacterial nutrient biosynthesis and antibiotic resistance are being revealed recently (Carfrae and Brown, 2023). These idiosyncratic connections offer an untapped opportunity for designing novel approaches to combat antibiotic resistant pathogens.

## Acknowledgements

Authors thank Nirma Education and Research Foundation (NERF), Ahmedabad for infrastructural support, and for providing stipend to NT. GG acknowledges scholarship from Government of Gujarat through their SHODH scheme.

