## Supplementary File for "Silversol^®^ (a colloidal nanosilver formulation) inhibits growth of antibiotic-resistant *Staphylococcus aureus* by disrupting its physiology in multiple ways"

**Table S1. Antibiogram of antibiotic-resistant *Staphylococcus aureus* generated through Kirby-Bauer Disc Diffusion assay**

| Antibiotic | Concentration (µg) | Interpretation |
|---|---|---|
| Penicillin G | 10 | S |
| Oxacillin | 1 | S |
| Erythromycin | 15 | R |
| Clindamycin | 2 | R |
| Linezoild | 30 | S |
| Co- Trimoxazole | 25 | S |
| Vancomycin | 30 | S |
| Tetracycline | 30 | S |
| Chloramphenicol | 30 | S |
| Gentamicin | 10 | S |
| Azithromycin | 15 | R |
| Ofloxacin | 5 | S |
| Methicillin | 5 | S |
| Amoxycillin/ Clavulanic acid | 30 | I |
| Clarithromycin | 15 | R |
| Ampicillin | 10 | S |
| Amikacin | 30 | S |
| Cephalothin | 30 | S |
| Novobiocin | 5 | S |
| Teicoplanin | 10 | S |

Antibiotic susceptibility profile of the bacterium was generated using the antibiotic discs- Icosa G-I Plus (HiMedia, Mumbai), through disc diffusion assay performed as per CLSI guidelines. The zones of inhibition were measured and the interpretation (S - sensitive, I - intermediate, R - resistant) was drawn as per zone size interpretative chart provided by the manufacturer.

**Table S2. Quantification of extracted RNA, library, and insert size**

| Sr. no. | Sample name | Quantification of extracted RNA | | | Library quantification and insert size analysis | |
|---|---|---|---|---|---|---|
| | | $OD_{260}/OD_{280}$ | $OD_{260}/OD_{230}$ | RIN value | ng/µL | Insert size |
| 1 | Control | 1.9 | 1.62 | 6.0 | 29 | 347 |
| 2 | Experimental | 1.64 | 1.7 | 4.2 | 22.6 | 341 |

**Table S3. Temperature profile for RT-PCR assay**

| Temperature (°C) | Time (s) | Remarks |
|---|---|---|
| **PCR cycles (45 cycles)** | | |
| 95 | 15 | Denaturation temperature |
| 59 | 60 | Annealing temperature |
| **Melt curve stage** | | |
| 95 | 15 | |
| 60 | 60 | |
| 95 | 15 | |

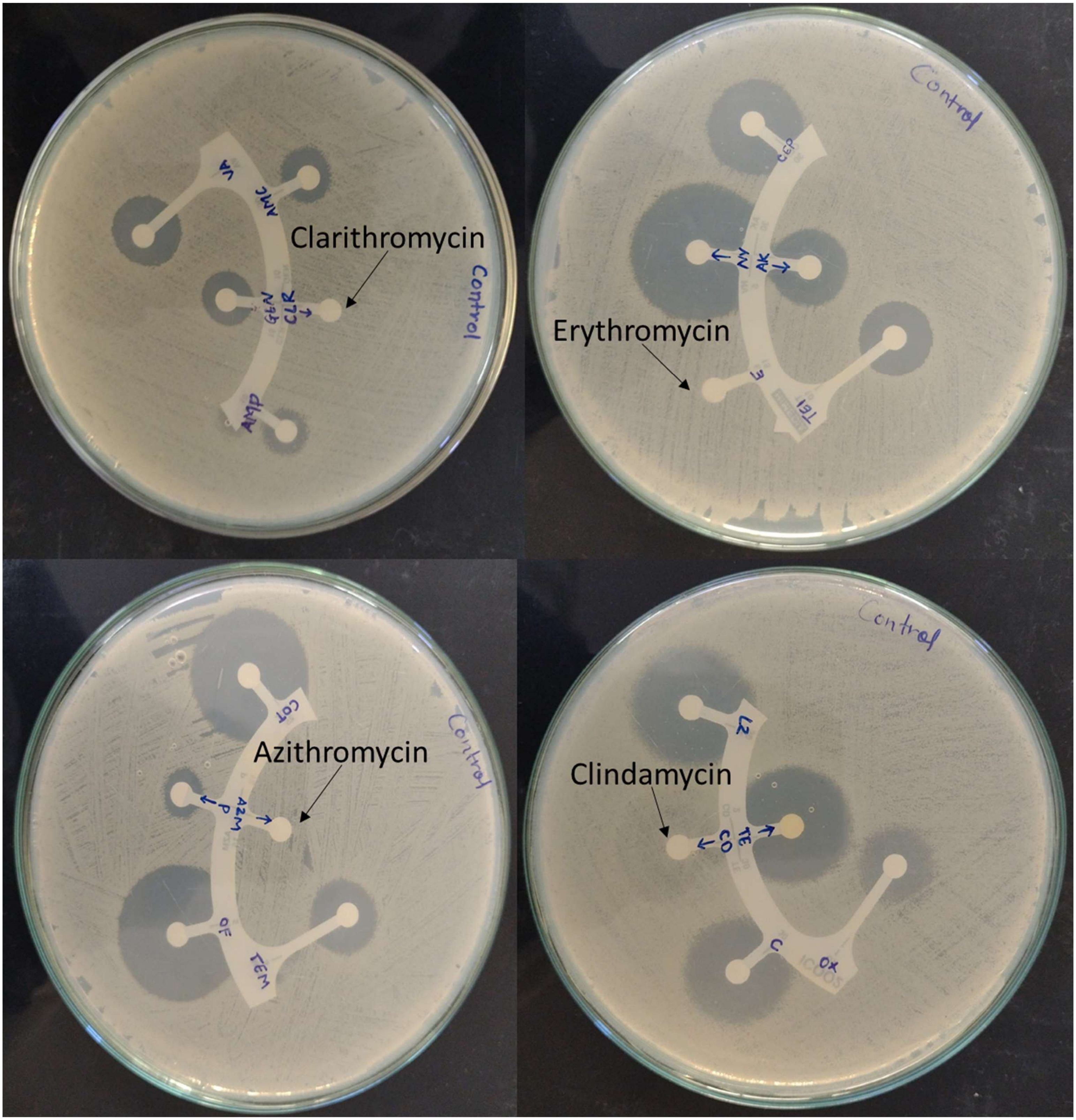


**Figure S1. Antibiogram of *S. aureus* generated through Kirby-Bauer Disc Diffusion assay**

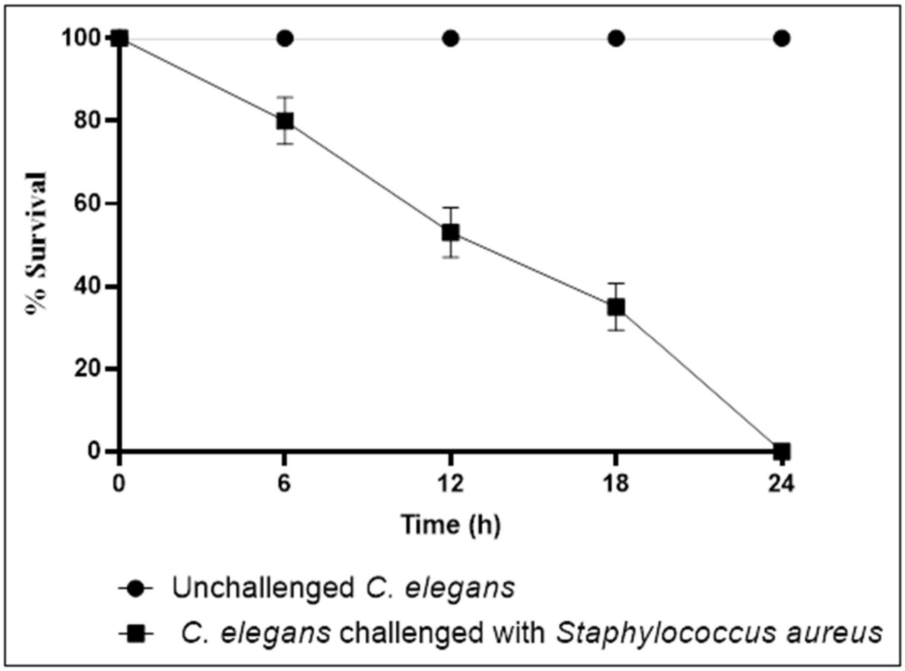


**Figure S2. Antibiotic-resistant strain of *S. aureus* used in our study could kill all the host worms within 24 h.** To confirm virulent nature of the *S. aureus* strain used by us, we challenged the model host *Caenorhabditis elegans* with *S. aureus* culture suspension ($OD_{764}$ = 1.50 grown in tryptone soy broth) in 24-well plate, and quantified worm survival through microscopic live-dead count.

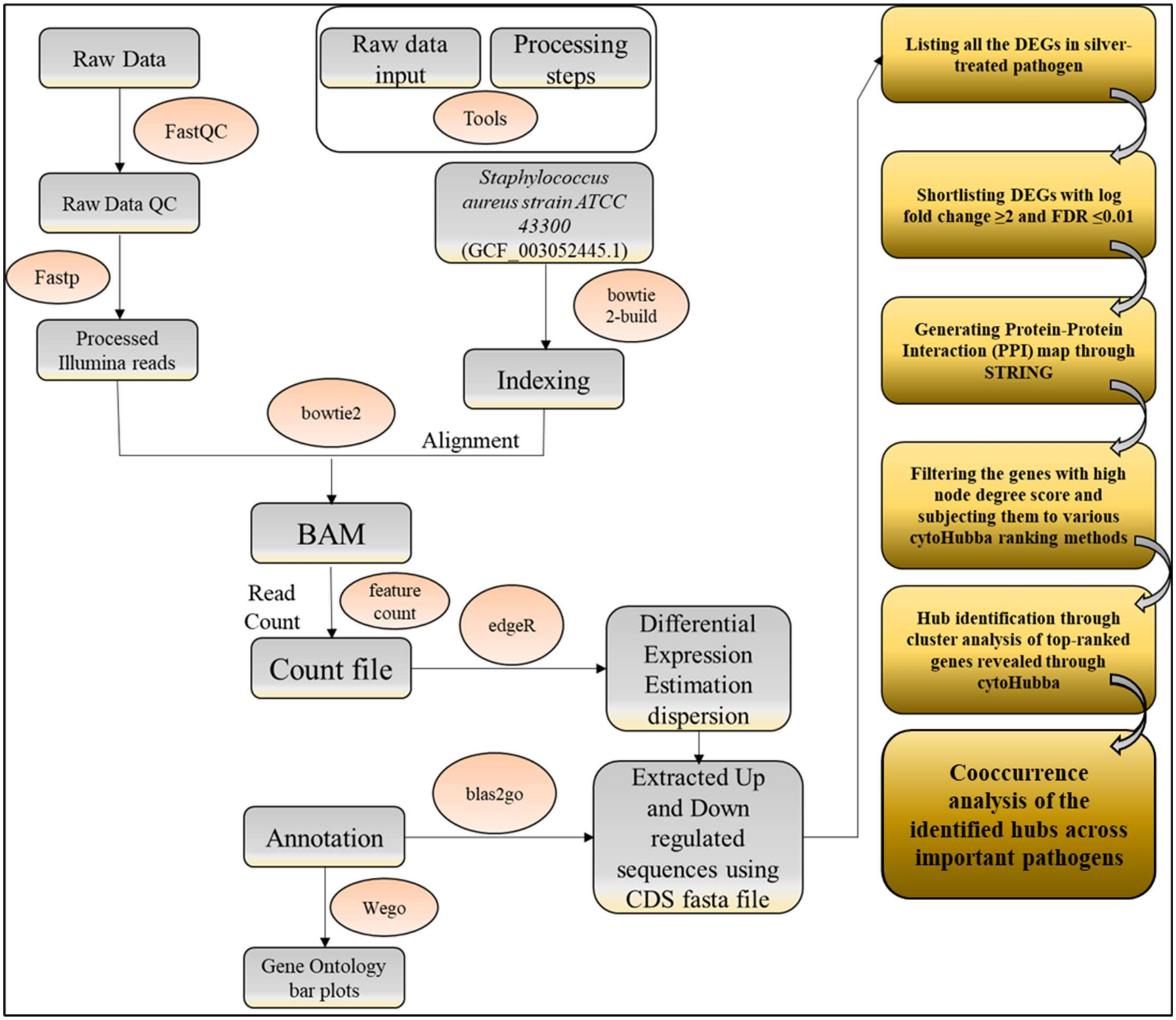


**Figure S3. A schematic presentation of the methodology/workflow employed for whole transcriptome and network analysis**

DEG: Differentially Expressed Genes

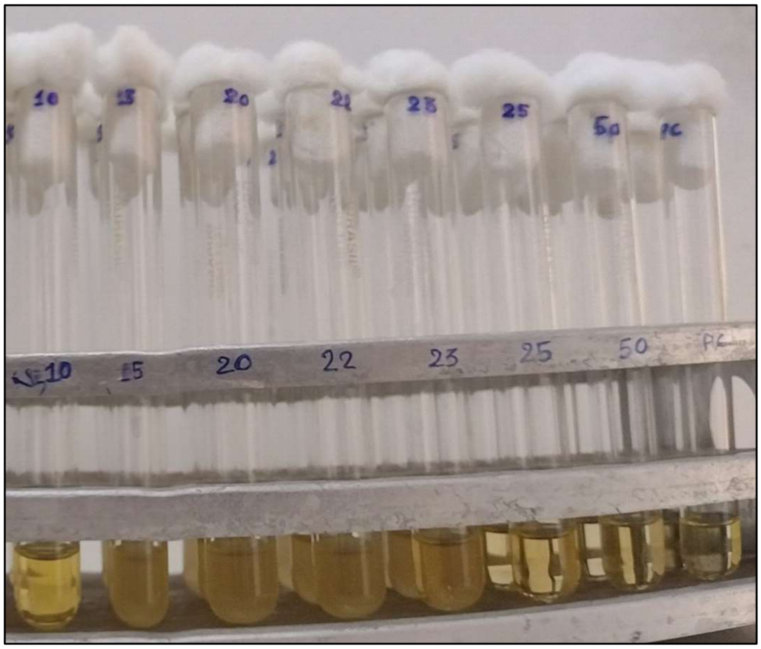


**Figure S4. Silversol's antibacterial effect against *S. aureus* does not follow a strictly linear dose-response pattern.** 15-23 ppm of Silversol were less effective than 10 ppm at inhibiting bacterial growth. The visible growth disappeared again 25 ppm onward. This image was captured after 24 h of incubation. Last tube corresponds to positive control, vancomycin (5 ppm).

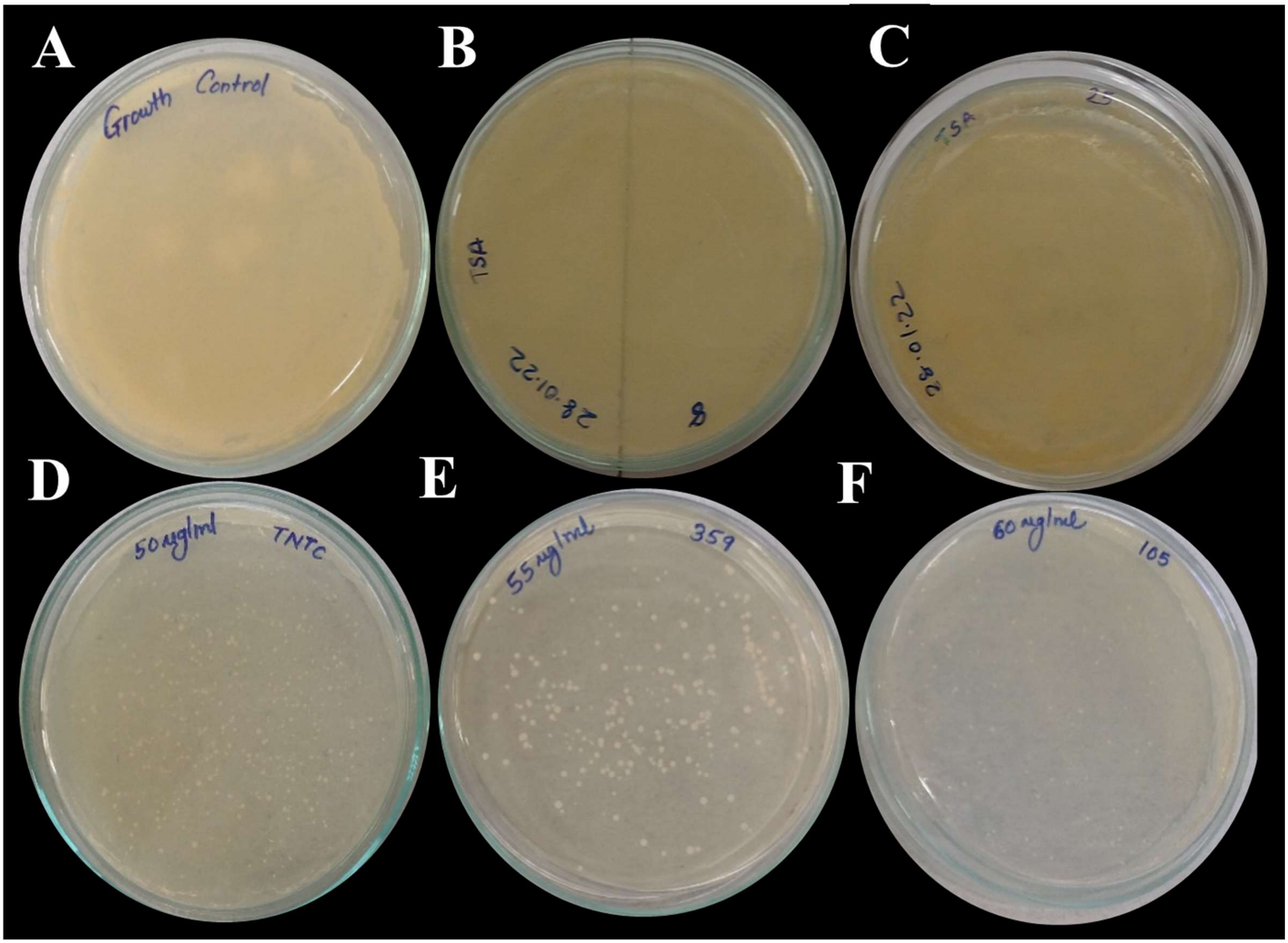


**Figure S5. Bacteriostatic and bactericidal effect of Silversol® against *S. aureus* at low and high concentrations respectively.** Cells grown in presence of Silversol® were subsequently plated onto silver-free tryptone soy agar. Cells sourced from 50-60 ppm Silversol-containing tubes resulted in few non-pigmented colonies, but these colonies took 72 hours post-inoculation to appear. Quantum of growth on these experimental plates was much lesser than on the control plate. However, cells sourced from lower Silversol concentration tubes could give rise to considerable growth indicating bacteriostatic effect. Images A-C were captured after 24 h of incubation, while D-F were captured after 72 h of incubation.

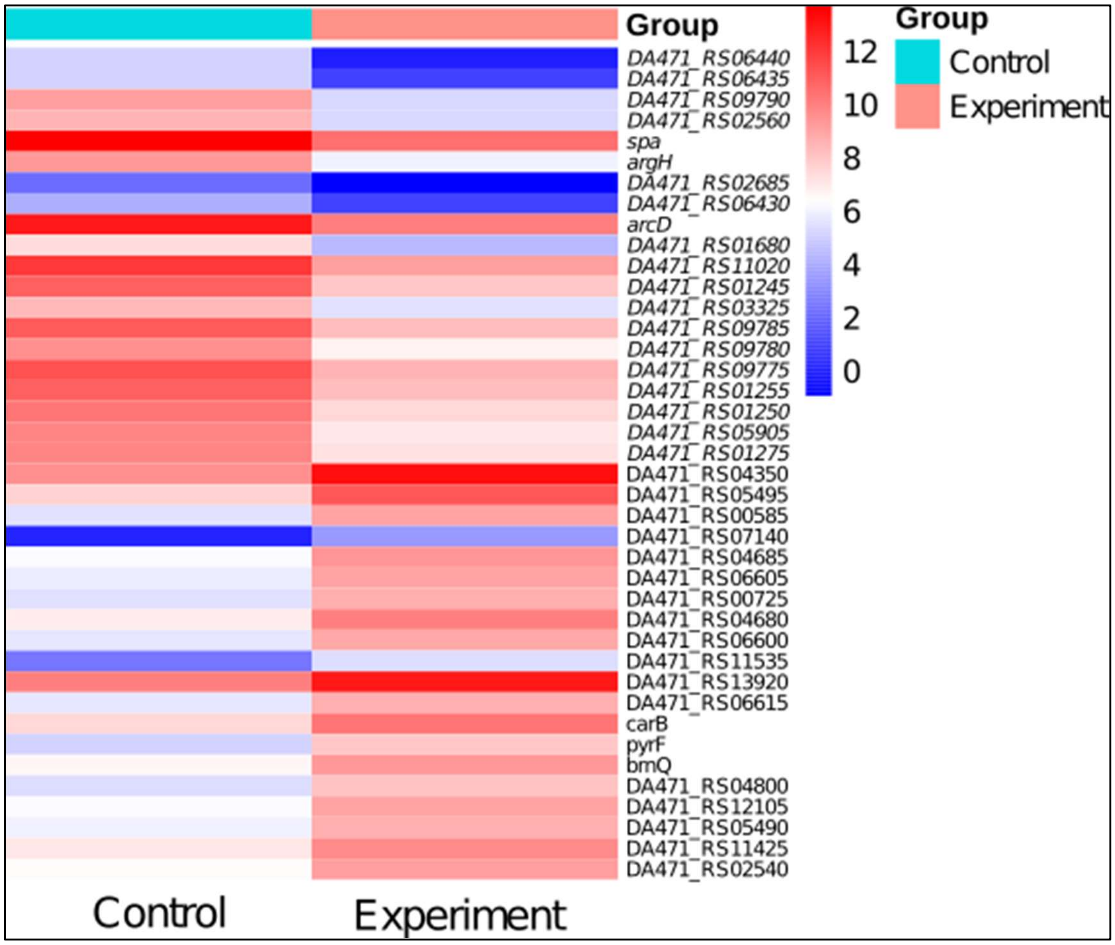


**Figure S6. Heat map of DEGs in Silversol®-exposed *S. aureus***

Heat map generated using the online software tool ClustVis (https://biit.cs.ut.ee/clust vis/) showing up-regulated and down-regulated genes with FDR<0.05 and log fold change '± 2'.

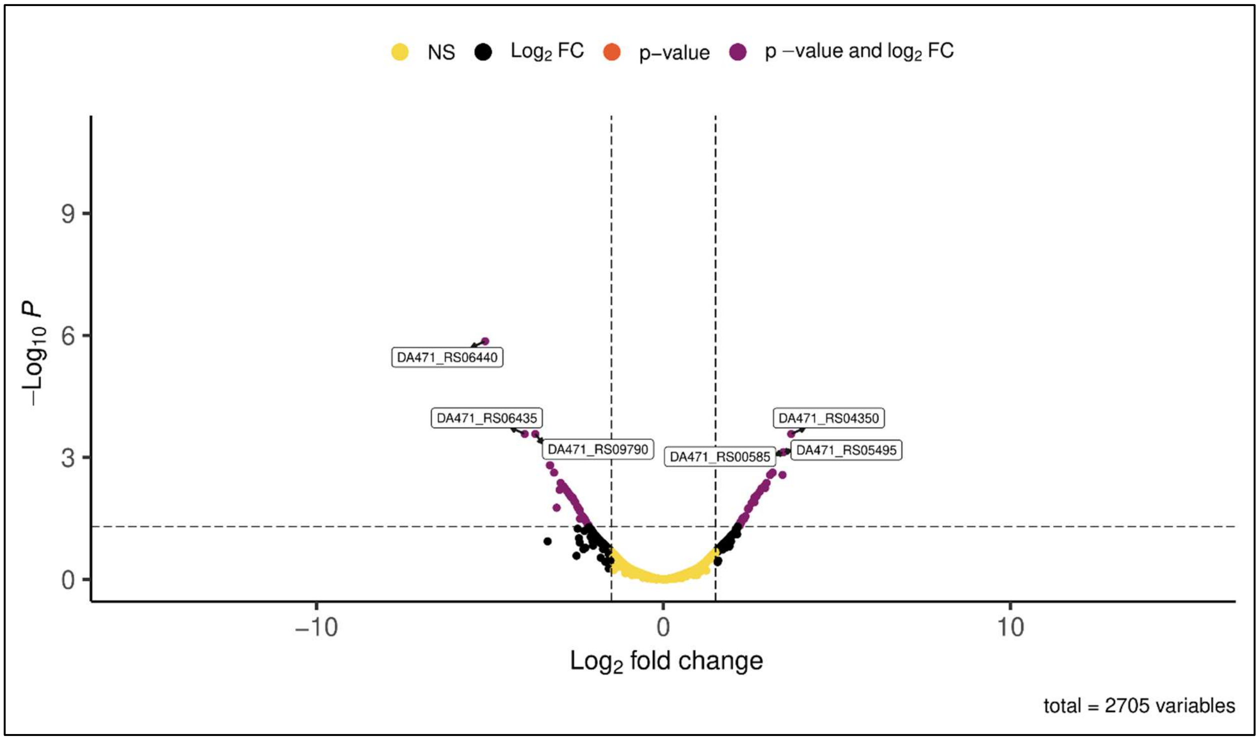


**Figure S7. Volcano Plot of experimental versus control samples**

Volcano plot of expressed genes of experimental culture compared to control culture. The y- axis illustrates −log 10 *p* values, and the x-axis corresponds to a log 2-fold change of gene expression between both cultures. The red points represent differently expressed genes satisfying the dual criteria of FDR≤0.05 and log fold change ≥2.
